# Stress-Induced Transcriptional Memory Accelerates Promoter-Proximal Pause-Release and Decelerates Termination over Mitotic Divisions

**DOI:** 10.1101/576959

**Authors:** Anniina Vihervaara, Dig Bijay Mahat, Samu V. Himanen, Malin A.H. Blom, John T. Lis, Lea Sistonen

## Abstract

Heat shock triggers an instant reprogramming of gene and enhancer transcription, but whether cells encode a memory to stress, at the level of nascent transcription, has remained unknown. Here, we measured transcriptional response to acute heat stress in unconditioned cells and in daughters of cells that had been exposed to a single or multiple heat shocks. Tracking RNA Polymerase II (Pol II) genome-wide at nucleotide-resolution revealed that cells precisely remember their transcriptional identity throughout stress, restoring Pol II distribution at gene bodies and enhancers upon recovery. However, single heat shock primed faster gene-induction in the daughter cells by increasing promoter-proximal Pol II pausing, and accelerating the pause-release. In repeatedly stressed cells, both basal and inducible transcription was refined, and pre-mRNA processing decelerated, which retained transcripts on chromatin and reduced recycling of the transcription machinery. These results mechanistically uncovered how the steps of pause-release and termination maintain transcriptional memory over mitosis.

**Highlights:**

-Cell type-specific transcription precisely recovers after heat-induced reprogramming
-Single heat shock primes genes for accelerated induction over mitotic divisions *via* increased promoter-proximal Pol II pausing and faster pause-release
-Multiple heat shocks refine basal and inducible transcription over mitotic divisions to support survival of the daughter cells
-Decelerated termination at active genes reduces recycling of Pol II to heat-activated promoters and enhancers
-HSF1 increases the rate of promoter-proximal pause-release *via* distal and proximal regulatory elements

## Introduction

Heat shock is one of the most potent triggers of transcription reprogramming, provoking an instant and genome-wide change in RNA synthesis from genes and enhancers (reviewed in Vihervaara *et al*., 2018). Upon heat shock, hundreds of genes are rapidly induced by potent *trans*-activators, such as Heat Shock Factor 1 (HSF1). When activated, HSF1 binds to Heat Shock Elements (HSEs) at architecturally primed promoters and enhancers (Rougvie and Lis, 1988; Rasmussen and Lis, 1993; Guertin and Lis, 2010; Vihervaara *et al*., 2013; 2017; Ray *et al*., 2019), and it can trigger the release of promoter-proximally paused Pol II into productive elongation (Duarte *et al*., 2016; Mahat *et al*., 2016). Concomitantly with the heat-induced escape of Pol II from the promoters of activated genes, thousands of genes are repressed *via* inhibition of the pause-release. This repression leads to the accumulation Pol II at promoter-proximal regions and clears transcription complexes from gene bodies (Mahat *et al*., 2016; Vihervaara *et al*., 2017). As a consequence of the genome-wide re-coordination of Pol II pause-release, heat-stressed cells promptly switch their transcription program to produce chaperones and protect cellular integrity.

Stress responses are evolutionarily conserved and robustly activated. Yet, severe stress can have long-lasting consequences for an individual (Guan *et al*., 2002; Sailaja *et al*., 2012) and cause physiological changes over several generations (Kaati *et al*., 2002; Wei *et al*., 2014; reviewed in Heard and Martienssen, 2014). On a cellular level, various stresses have been associated with long-term changes in the chromatin state (Guan *et al*., 2002; Tetievsky and Horowitch, 2010; Sailaja *et al*., 2012; D’Urso *et al*., 2016; Lämke *et al*., 2016; reviewed in D’Urso and Brickner, 2017), and shown to confer thermotolerance against protein misfolding *via* increased chaperone expression (Gerner and Schneider, 1975; Maytin *et al*., 1990; Yost and Lindquist, 1991). To date, however, stress-induced long-term changes in gene expression have been investigated from steady-state RNA and protein expression, which do not quantify processes of nascent transcription or reveal mechanistic control of Pol II. Besides chaperone expression, acquired stress resistance is commonly assessed as improved cell viability. Since apoptosis is the final stage of failed proteostasis and involves radical changes in cellular physiology, evaded cell death is not an appropriate stage to track control of nascent transcription that underlies the onset of stress resistance. Consequently, we do not yet know whether cells restore or adjust their program of RNA synthesis when recovering from heat stress, and whether nascent transcription profile can encode a memory to previously encountered proteotoxicity.

In this study, we used heat shock to provoke a genome-wide change in gene and enhancer transcription, and asked whether proteotoxic stress reprograms nascent transcription and transcriptional responsiveness over mitotic divisions. To map RNA synthesis at nucleotide resolution, we used Precision Run-On sequencing (PRO-seq) that tracks incorporation of a single biotinylated nucleotide at the active site of each nascent transcript (Kwak *et al*., 2013). Consequently, PRO-seq provides genome-wide maps of transcribing Pol II complexes at genes and enhancers, and identifies regulatory decisions at high fidelity and spatio-temporal resolution (reviewed in Wissink *et al*., invited manuscript under review). As cell models, we used Mouse Embryonic Fibroblasts (MEFs) and human K562 erythroleukemia cells that reprogram transcription upon heat shock with highly similar mechanisms (Mahat *et al*., 2016; Vihervaara *et al*., 2017). We found that transcriptional reprogramming by heat shock is followed by precise restoration of basal cell type-specific transcription program of genes and enhancers. The robustly restored transcription profile allowed addressing mechanisms that convey memory to heat shock, without concurrently triggering a change in the cell’s transcriptional identity or function: A single heat shock primed a subset of genes for instant induction in the daughter cells by increasing promoter-proximal Pol II pausing and by accelerating the pause-release upon an additional heat shock. Repeatedly stressed cells, in turn, refined transcription program over mitotic divisions by decreasing the synthesis of genes for protein production and by increasing expression of genes for pro-survival. Moreover, repeated heat shocks prolonged Pol II residency at the Cleavage and Polyadenylation Site (CPS) of active genes, concurrently reducing transcript cleavage and recycling of Pol II. These results revealed stress-induced control of Pol II that establishes and maintains transcriptional memory over mitotic divisions.

## Results

### Gene and Enhancer Transcription Is Precisely Restored after an Acute Heat Shock

To address whether heat-induced reprogramming of RNA synthesis is followed by restoration or readjustment of transcription, we measured nascent RNA synthesis in MEFs upon a 1-hour heat shock, and after a 4-hour or a 48-hour recovery (Figure 1). DNA staining and Fluorescence Associated Cell Sorting (FACS) verified that MEFs proliferated over the acute heat shock and also following recovery, and they did not undergo cell cycle arrest or apoptosis (Figure S1A). For accurate normalization of PRO-seq data, we used whole-genome spike-in from *Drosophila* S2 cells (Figure S1B, see Materials and Methods for details), and pooled biological replicates for data analyses after validating their high correlation (Figure S1C). As previously reported (Mahat *et al*., 2016; Vihervaara *et al*., 2017), a single heat shock caused a robust induction of hundreds and repression of thousands of genes, simultaneously accumulating Pol II at divergently transcribed distal Transcription Regulatory Elements (dTREs), hereon called enhancers (Figure 1A). During a 4-hour recovery, the genome-wide profile of gene and enhancer transcription was precisely restored to the level observed prior to the heat shock (Figure 1A-B), which demarcates high plasticity and accurate recovery of the transcription program. Despite the full recovery of gene body transcription, certain promoter-proximal regions gained new pause sites, as exemplified by *Heat Shock Protein H1* (*Hsph1 alias Hsp110*) gene (Figure 1B). Moreover, a subset of heat-responsive genes either gained (Figure S1D) or retained (Figure S1E) high Pol II density at a single pause site during the recovery. Consequently, the genome-wide average of Pol II at the pause region remained elevated, even when measured in the daughter cells, 48 hours after the heat exposure (Figure 1C).

**Figure 1.**
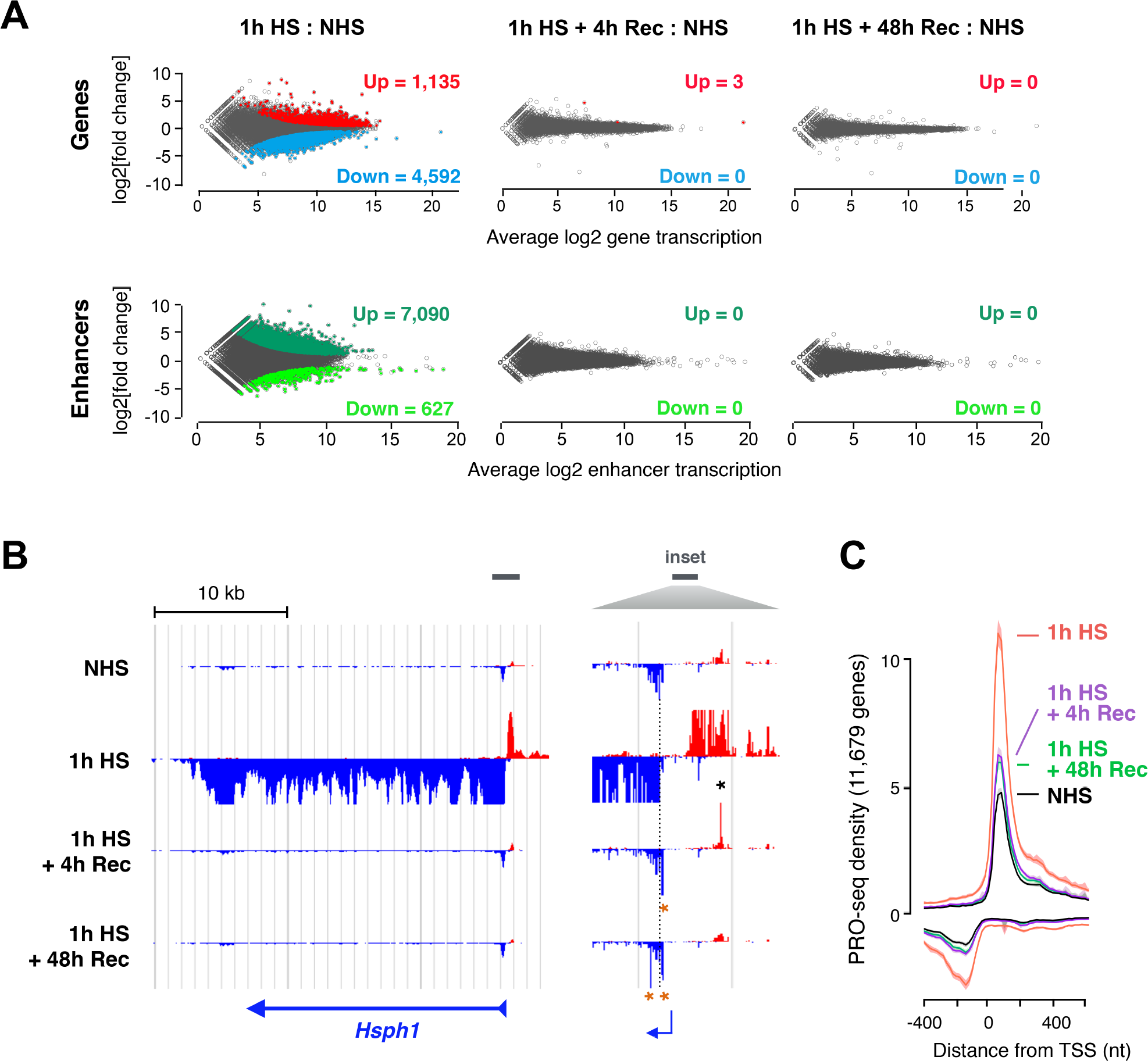
Transcription of genes and enhancers precisely recovers after heat-induced reprogramming. **A)** DESeq2-analysis of differential gene and enhancer transcription in mouse embryonic fibroblasts (MEFs). Up and Down denote a statistically significant increase or decrease, respectively, in Pol II density at gene bodies (upper panels) and enhancers (lower panels) upon heat shock and recovery, as compared to optimal growth conditions. **B)** Transcriptional profile of a heat-induced *Hsphl* gene in the non-heat-shock condition, upon 1 h of heat shock, and upon recovery from a 1-h heat shock. Inset depicts promoter-proximal region. The dashed line indicates highest Pol II pausing density in non-heat-shocked cells, and asterisks denote prominent Pol II pausing on sense (orange) and anti­ sense (black) strand during recovery. **C)** Average promoter-proximal pausing measured at all transcribed genes. Shaded area indicates 12.5 to 87.5% confidence interval. HS: heat shock; NHS: non-heat shock; Rec: recovery.

### Heat Shock Primes Accelerated Gene Induction over Mitotic Divisions

Since individual genes and whole transcription programs can be coordinated at the promoter-proximal pause-release (Rougvie and Lis, 1988; Boettiger and Levine, 2009; Mahat *et al*., 2016; Vihervaara *et al*., 2017), we addressed whether the changed pausing pattern in daughter cells alters genes’ heat responsiveness. We preconditioned MEFs with a single 1-hour heat shock, and allowed a 48-hour recovery before measuring transcription kinetics provoked by an additional heat shock. To capture instant and sustained changes in heat-induced transcription, we conducted PRO-seq upon 0, 12.5, 25, and 40 minutes of heat shock, and compared transcriptional stress response between unconditioned and preconditioned cells (Figure 2A; Figure S1F-H). Certain heat-induced genes, such as *Ubc* that encodes polyubiquitin (Figure 2B), gained promoter-proximal Pol II pausing during preconditioning (0-minute inset in Figure 2B), and released the paused Pol II faster into elongation upon heat shock (12.5-minute inset in Figure 2B). By 40 minutes of heat shock, unconditioned cells also had gained high *Ubc* transcription and efficient Pol II pause-release (Figure 2B), suggesting that preconditioning accelerated the onset, rather than changed the overall level of transcription. The faster activation was also demonstrated by heat-induced waves of Pol II progression: For example at *Vinculin* (*Vcl*) gene, the advancing wave of transcription had proceeded farther during the 12.5-minute heat shock in preconditioned than in unconditioned cells (Figure 2C). This transiently activated gene had returned to its basal expression by 40 minutes of heat shock, regardless of the preconditioning (Figure 2C). Importantly, *Vcl* did not show higher pausing in preconditioned cells (0-minute inset of Figure 2C), but the altered usage of promoter-proximal pause sites and the lower pausing signal upon heat stress (insets of Figure 2C) indicated faster kinetics of Pol II when proceeding from its promoter-proximal pause to productive elongation.

**Figure 2.**
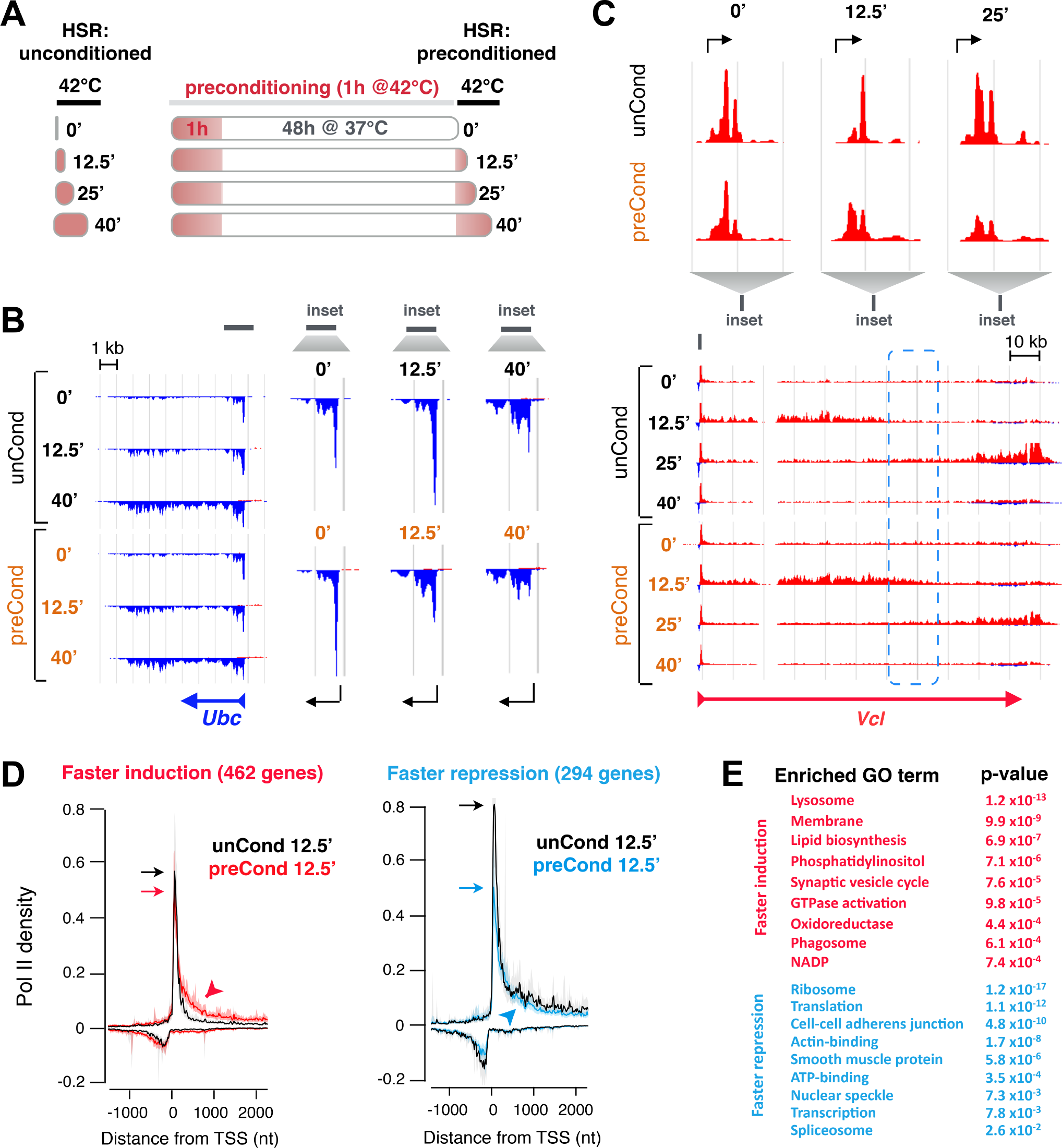
Single heat shock primes accelerated gene-induction. **A)** Experimental setup for measuring transcription kinetics in unconditioned (unCond) and single heat shock-conditioned (preCond) mouse embryonic fibroblasts (MEFs). Transcription was analyzed upon a 0, 12.5, 25 or 40-min heat shock in unconditioned cells (left panel), and in cells that were preconditioned with a 1-h heat shock and 48-h recovery (right panel). **B)** Transcriptional profile of *Ubc* prior to and upon heat shock in unconditioned and preconditioned cells. The insets show Pol II density in promoter-proximal region in unconditioned (upper panels) and preconditioned (lower panels) cells. **C)** Heat-induced wave of transcription along *Vcl* gene. The blue dashed region indicates an advancing wave of transcription that has proceeded farther in preconditioned than in unconditioned cells upon 12.5 min heat shock. The insets show promoter-proximal Pol II density in unconditioned (upper panels) and preconditioned (lower panels) cells. **D)** Average intensity of transcription upon a 12.5-min heat shock at the promoter-proximal region of genes that gain faster heat-induction (left panel) or heat-repression (right panel) by preconditioning. The highest pause-density in each condition is indicated with an arrow, the signal downstream of the pause is denoted with an arrowhead. **E)** Enriched gene ontology terms among genes that gain faster heat-induction (red) or heat-repression (blue) by preconditioning.

### Faster Pause-Release Accelerates Gene Induction in Preconditioned Cells

To examine whether the faster release of Pol II into productive elongation served as a general mechanism for accelerated induction, we compared transcription programs upon heat shock between unconditioned and preconditioned cells. The instant 12.5-minute heat shock showed the highest variation in gene expression due to preconditioning (Figure S2A), which supports observations from individual genes that preconditioning with a single heat shock primarily affected the onset of the transcriptional response (Figure 2B-C; Figure S2B). Noteworthy is that these changes in nascent RNA synthesis, as measured at the steps of initiation, pause-release and ongoing elongation, can only be detected with assays that directly track engaged Pol II complexes (reviewed Wissink *et al*., invited manuscript under review).

Of all heat-induced genes, 426 displayed significantly higher level of elongating Pol II in preconditioned cells upon the instant 12.5-minute heat shock (Figure S2A), which indicates accelerated heat induction. Simultaneously, 294 heat-repressed genes showed deeper decline in transcription due to preconditioning (Figure S2A). On average, genes with accelerated induction contained similar or slightly lower density of the engaged Pol II at promoter-proximal regions in preconditioned cells than in unconditioned cells upon 12.5 minutes of heat shock (Figure 2D). However, more Pol II had escaped into gene body transcription in preconditioned cells (Figure 2D; Figure S2A). This pattern of engaged Pol II molecules demonstrated faster progression of Pol II from initiation into elongation in the daughters of heat-shocked cells. Importantly, we did not detect preconditioning to accelerate heat responsiveness of genes that were instantly activated already in unconditioned cells (Figure S2C). Neither was kinetics of the classical heat shock genes changed (Figure S2D). Consequently, preconditioning with a single heat shock did not globally reduce response time, but primed an additional set of genes for a rapid heat-induction.

### Accelerated Induction of Quality Control Genes Co-occurs with Deeper Repression of Genes for Protein Production

Genes that gained accelerated heat induction by preconditioning were enriched for lysosomal, autophagocytosis and membrane-associated functions (Figure 2E), suggesting their role in acquired stress resistance. Genes with faster repression, instead, contained classical heat-repressed genes (Mahat *et al*., 2016; Vihervaara *et al*., 2017) that are highly expressed under non-stress conditions and involved in transcription, translation, and cell-cell junctions (Figure 2E). Across their promoter-proximal (Figure 2D) and gene body (Figure S2A) regions, genes with faster repression showed modestly lower Pol II density in heat-shocked preconditioned cells, which demonstrates fewer Pol II molecules engaging into their transcription. This reduced initiation could be indirectly caused by limited Pol II availability, as a larger fraction of Pol II molecules were instantly engaged into transcription of heat-activated genes. Taken together, our results in singly heat shock-conditioned MEFs demonstrated that cells precisely remembered their transcriptional identity throughout an acute heat shock, but that they also simultaneously pre-wired an additional set of genes for a more rapid induction in the daughter cells.

### Multiple Heat Shocks Reprogram Basal Transcription in Cancer Cells

Pathophysiological stresses, such as those encountered during cancer progression and neurodegeneration, are often sustained or repeated. To be able to investigate how repeatedly encountered stress affects gene and enhancer transcription, we moved from stress-sensitive MEFs to human K562 erythroleukemia cells. K562 cells are a patient-derived cancer cell line (Lozzio and Lozzio, 1975; Koeffler and Kolde, 1980), known to tolerate extended heat treatments and develop thermotolerance (Mivecchi, 1989; Vihervaara *et al*., 2013). We preconditioned K562 cells with a series of heat shocks, giving a total nine 1-hour heat treatments during three consecutive days, and allowed for a 48-hour recovery before measuring transcriptional response to an additional single heat shock (Figure 3A). It is important to note that K562 cells proliferated throughout the six days of preconditioning, recovery and additional heat shock (Figure S3A) without showing signs of apoptosis or increased polyploidy (Figure S3B). The gene and enhancer transcription in unconditioned and preconditioned cells was measured using PRO-seq with whole-genome spike-in normalization (Figure S3C-D).

**Figure 3.**
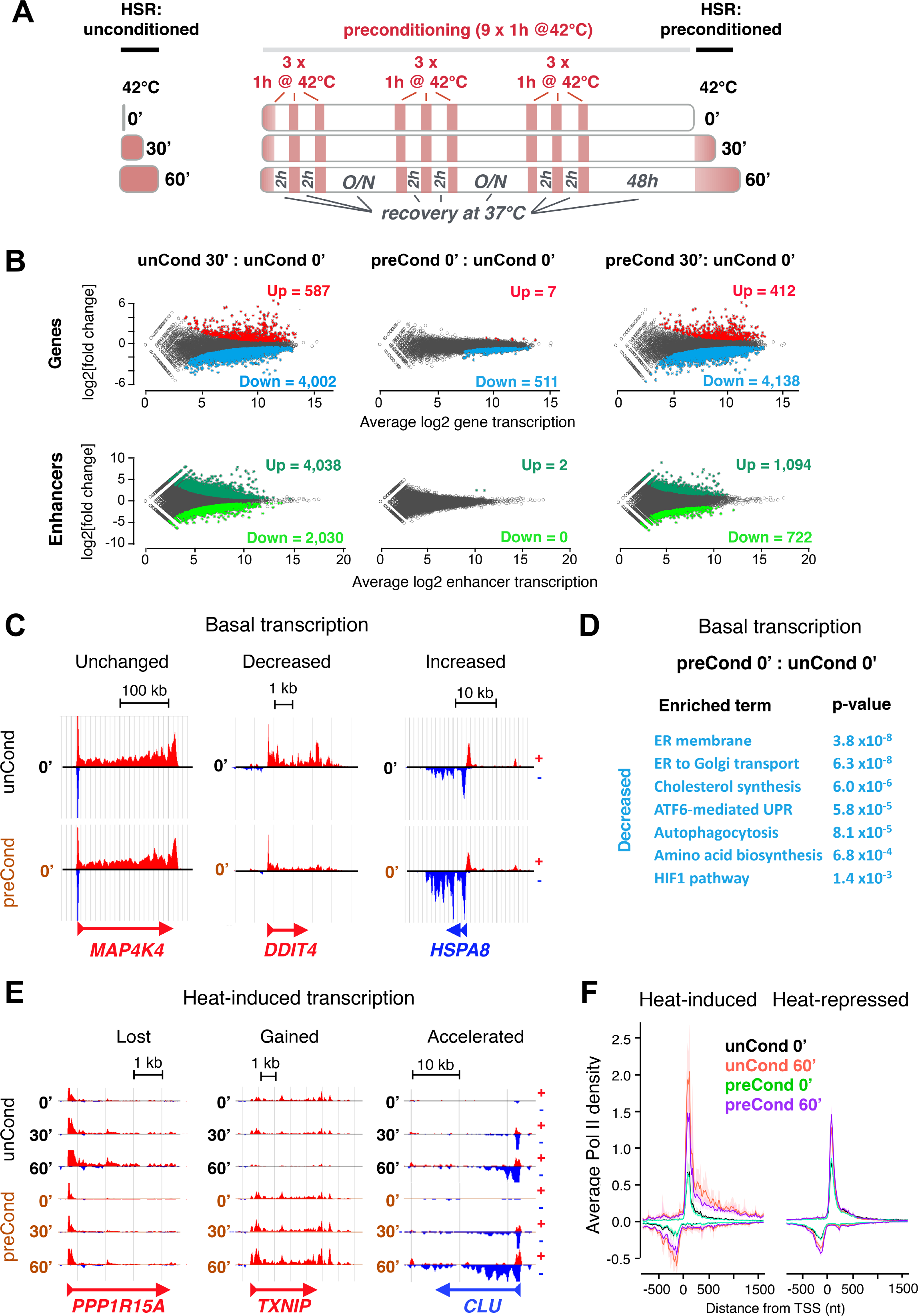
Repeated heat shocks refine both basal and inducible transcription in human K562 cells. **A)** Experimental setup for preconditioning human K562 cells with multiple heat shocks: Genome-wide nascent transcription was measured upon a 0, 30 or 60-min heat shock in unconditioned cells (left panel), and in cells that were pre-exposed to nine 1-h heat shocks during three consecutive days, and allowed to recover for 48 h prior to an additional 1-h heat shock (right panel). **B)** DESeq2-analysis of differential gene and enhancer transcription. Up and Down denote numbers of genes (upper panels) and enhancers (lower panels) with significantly heat shock-induced or repressed Pol II density, respectively, as measured against non-stressed cells (unCond 0’). **C)** Transcriptional profiles of genes with unchanged (left panel), decreased (middle panel) or increased (right panel) basal transcription due to preconditioning. **D)** Enriched gene ontology terms among genes that show decreased basal transcription in preconditioned cells. **E)** Transcriptional profiles of genes that had lost (left panel), gained (middle panel), or accelerated (right panel) heat-induction as a result of preconditioning. **F)** Average Pol II density at the promoter-proximal regions of heat-induced (left) and heat-repressed (right) genes in the indicated heat shock conditions.

In K562 cells, a single heat shock induces hundreds and represses thousands of genes, simultaneously increasing Pol II density at transcribed enhancers (Figure 3A-B; Vihervaara *et al*., 2017). During the 48-hour recovery from preconditioning, the vast majority of genes and virtually every enhancer restored their transcription (Figure 3B; Figure S3E), which indicates robust recovery of cell type-specific RNA synthesis even after multiple heat exposures. Preconditioning with several heat shocks, however, changed basal transcription of a subset of genes, causing higher synthesis of only seven, and reduced synthesis of over 500 genes (Figure 3B-C). The most prominent increase in basal transcription was detected for *HSPA8* (Figure 3C) that encodes HSP70 cognate (HSC70), a constitutively expressed chaperone known to safeguard protein homeostasis (Ignolia and Craig, 1982; Kampinga *et al*., 2009). In contrast, genes with repressed basal transcription encoded regulators of protein production and maturation, including DDIT3, HSPA5, XBP1 and HSP90AB1 of the Endoplasmic Reticulum (ER) and Golgi pathway (Figure 3D, Supplementary Dataset 1). The uncompromised proliferation (Figure S1A-B) and unchanged expression of cell cycle regulators (Figure S3E), together with the adapted transcription program to maintain homeostasis (Figure 3D, Supplementary Dataset 1), indicated that K562 cells proliferated under repeatedly encountered protein-damaging conditions, and carried stress-induced transcription changes over cell divisions to their daughter cells.

### Repeated Stress Re-Wires Heat-Inducibility

Heat-responsiveness was restored for the vast majority of genes in preconditioned K562 cells (Figure 3B), but a subset of genes lost, gained or accelerated heat induction due to preconditioning (Figure 3E). One of the genes that had lost heat induction encodes Protein Phosphatase 1 Regulatory subunit 15A (PPP1R15A *alias* GADD34; Figure 3E) that counteracts stress-induced inhibition of translation by dephosphorylating eukaryote Initiation Factor 2 alpha (eIF2α; Harding *et al*., 2009; Walter and Ron, 2011). In support of inhibited protein synthesis in preconditioned cells, the few genes that had gained heat induction included proteins with function in cell survival and growth arrest (Supplementary Dataset 1). Albeit certain genes gained accelerated heat induction (Figure 3E), we did not detect increased Pol II density at the promoter-proximal regions (Figure S3F), as was detected in MEFs (Figure 1C) after a 48-hour recovery from stress. On the contrary, 5’-sequences of heat-induced genes gained less Pol II in preconditioned K562 cells upon heat shock (Figure 3F). The differences in Pol II entry into heat-activated genes indicated either cell-type specificity, or more likely, diverging strategies to re-wire transcriptional responsiveness in cells that were exposed to either a single or multiple heat shocks.

### Repeated Heat Shocks Change Pol II Progression through Heat-Induced Genes

To mechanistically understand why the promoter-proximal regions of heat-induced genes gained low Pol II density in preconditioned K562 cells (Figure 3F), we measured recruitment of HSF1 to the promoters of classical heat shock genes *HSPA1A* and *HSPH1*, and analyzed their nascent RNA synthesis and expression of mature mRNAs (Figure 4A; Figure S4A). Strikingly, HSF1-binding to the promoters of *HSPA1A* and *HSPH1* was similar in unconditioned and preconditioned cells (Figure 4A; Figure S4A), indicating uncompromised capacity of HSF1 to bind to its target regulatory elements. In contrast, RNA synthesis of *HSPA1A* and *HSPH1* was blunted, and the levels of their corresponding mature mRNAs were lower in preconditioned cells (Figure 4A; Figure S4A). These results demonstrated that mechanisms beyond HSF1 binding caused reduced expression of heat-responsive genes in preconditioned cells. PRO-seq profiles of nascent transcription suggested compromised Pol II progression to underlie the lower RNA expression of heat-induced genes. Reduced Pol II density at the promoter-proximal regions was accompanied with a locally confined increase in engaged Pol II downstream of the CPS, and this pattern of Pol II progression was detected at *HSPH1* (Figure 4B), as well as along highly heat-induced genes in general (Figure 4C).

**Figure 4.**
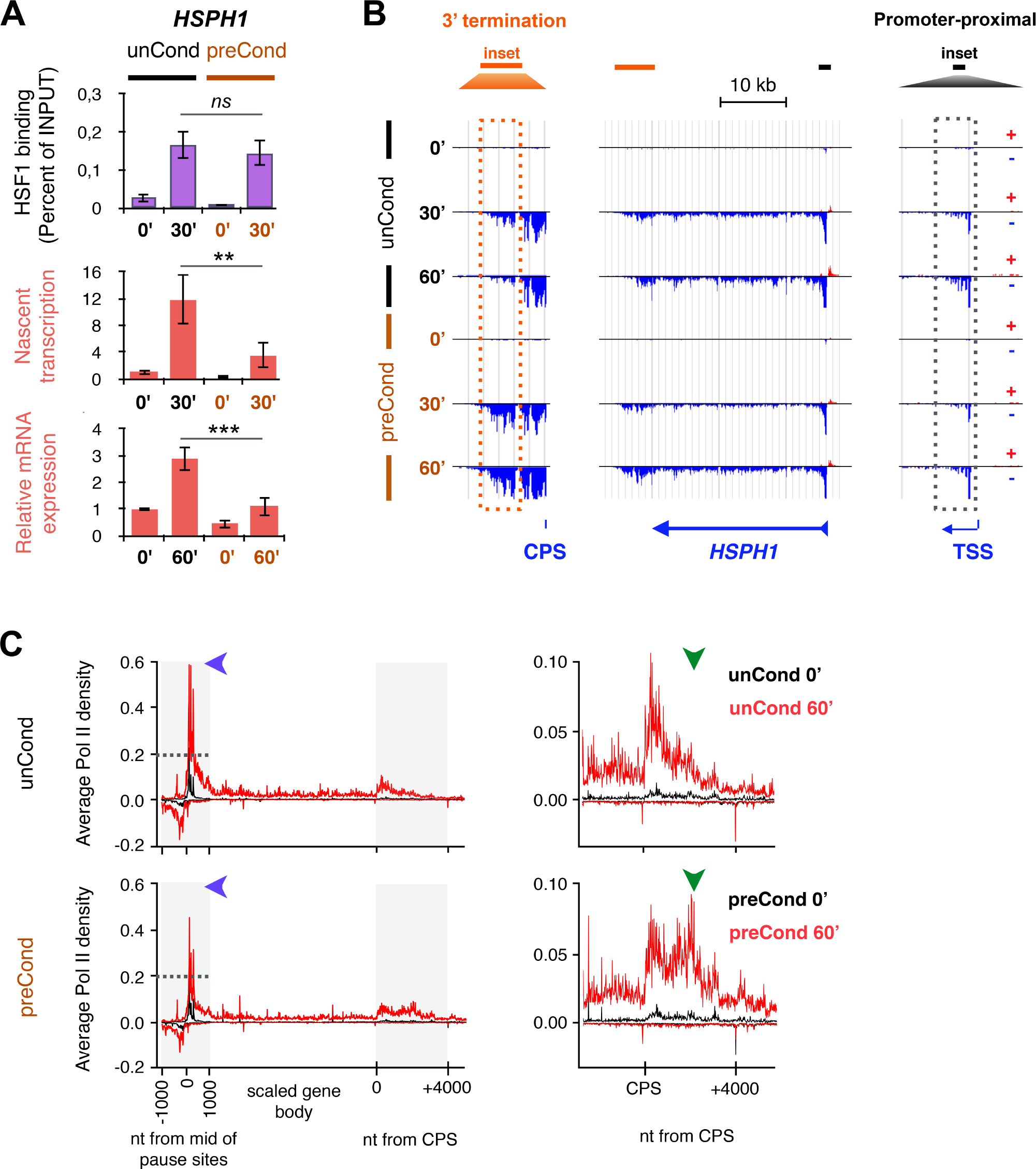
HSF1-binding to target gene promoters is not compromised, but the subsequent nascent transcription and mRNA expression are blunted in preconditioned cells. **A)** HSF1-binding intensity to the *HSPHJ* promoter (uppermost panel), nascent transcription of *HSPHJ* as measured from the first intron, +1055 to +1116 from the TSS, (middle panel), and relative level of polyA-containing *HSPHJ* mRNA (bottom panel) in unconditioned and preconditioned K562 cells. **B)** Nucleotide-resolution profile of engaged Pol II along *HSPHJ* showing reduced Pol II density at the promoter-proximal region and increased Pol II density downstream of the Cleavage and Polyadenylation Site (CPS). **C)** Average density of the engaged Pol II along highly heat-induced genes in unconditioned (upper panel) and preconditioned (lower panel) cells. Blue arrowheads indicate the pausing signal in unconditioned cells, and green arrowheads show the site of increased Pol II density in the 3’ ends of preconditioned cells.

### HSF1-Dependent eRNA Transcription Is Diminished in Preconditioned K562 Cells

To quantify HSF1-dependent gene and enhancer transcription genome-wide, we tracked nascent transcription in HSF1-depleted K562 cells (Figure 5A). In total, 227 genes and 496 enhancers were HSF1-dependently heat-induced both in unconditioned and preconditioned cells (Figure S4B). This data demonstrates that HSF1 *trans*-activates both genes and enhancers, and that it is indispensable for the heat-induced regulatory element activation regardless of the preconditioning (Figure 5B-E; Figure S4B-C). The ability of HSF1 to *trans*-activate genes from enhancers was evident for genes such as *Tax1 Binding Protein 1* (*TAX1BP1*). At this locus, HSF1 only bound to a divergently transcribed enhancer 4.5 kb upstream of the promoter (Figure 5C-D). However, HSF1 was indispensable besides for the heat-induced enhancer RNA (eRNA) transcription, also for the release of Pol II from the *TAX1BP1* promoter into productive elongation (Figure 5C-E).

**Figure 5.**
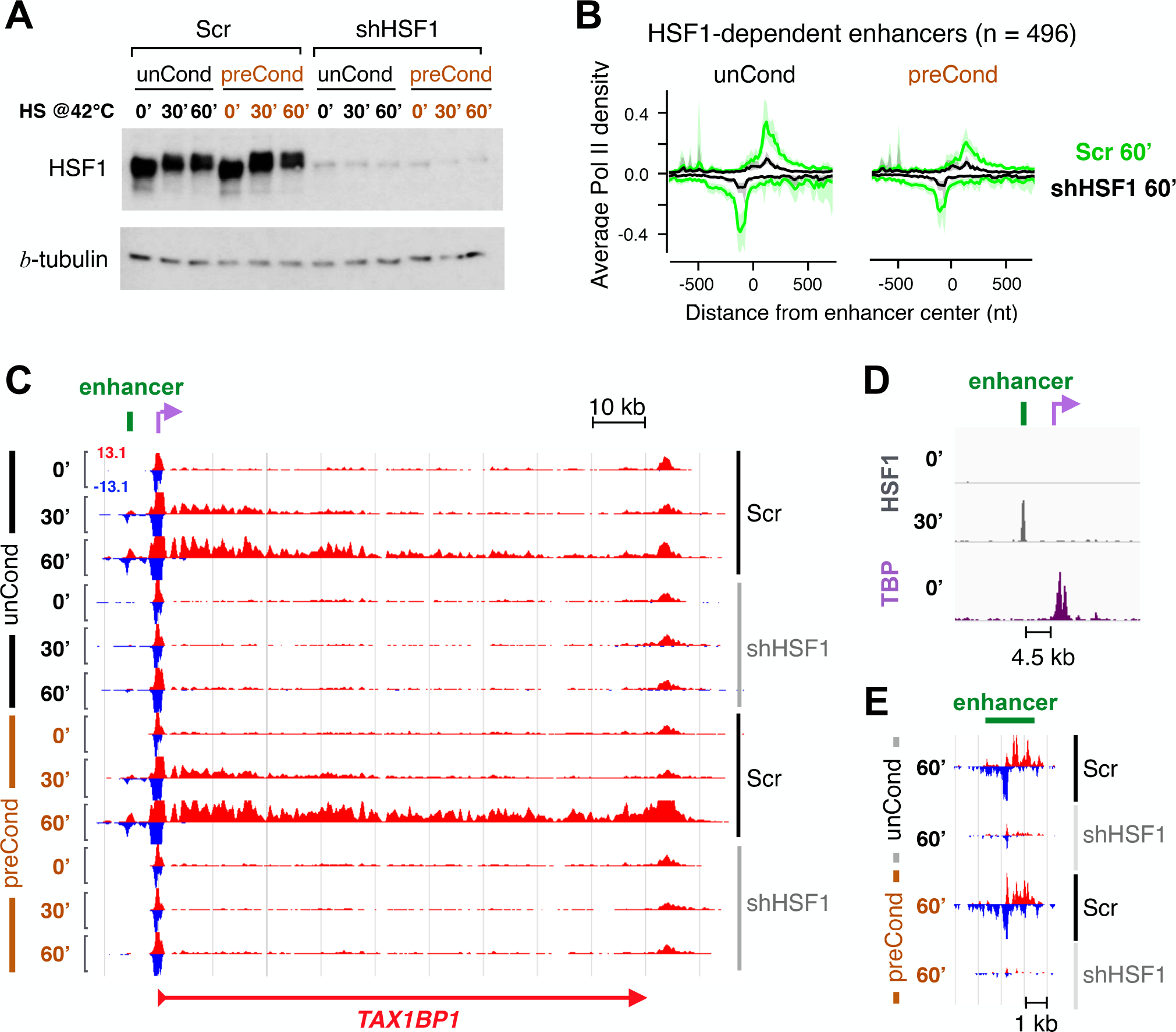
HSF1 *trans*-activates genes *via* enhancers, and is indispensable for heat-activation of gene and enhancer transcription in unconditioned and preconditioned cells. **A)** HSF1 protein expression in scrambled-transfected (Ser) and HSF1-depleted (shHSF1) K562 cells. Beta-tubulin serves as a loading control. **B)** Average Pol II density at HSFI­ dependently heat-induced enhancers in the presence (Ser) and absence (shHSF1) of HSFI. **C-E)** HSF1 drives heat­ induced transcription of *TAXIBPI* gene *via* an upstream enhancer. **C)** Transcription of *TAXIBP1* and its upstream enhancer in the presence (Ser) and absence (shHSF1) of HSFI in unconditioned and preconditioned cells. **D)** Inset of *TAXIBPI* enhancer (green bar) and TSS (purple arrow), showing heat-induced HSF1 binding (gray) to the enhancer, and IBP localization to the promoter (purple). **E)** Inset showing enhancer transcription in the presence and absence of HSF1 in unconditioned and preconditioned cells upon a 60-min heat shock. IBP: TATA box Binding Protein. ChIP-seq data for IBP was obtained from ENCODE (Consortium EP, 2011) and for HSFI from Vihervaara *et al.* (2013).

HSF1 can release the promoter-paused Pol II *via* proximal (Duarte *et al*., 2016; Mahat *et al*., 2016) and distal (Figure 5C-E) regulatory elements, but the mechanisms by which HSF1 activates enhancers remain unknown. Enhancers are also regulated at the step of Pol II pause-release (Henriques *et al*., 2018), but this pause is short-lived and we have not detected Pol II pausing to prime enhancers for heat activation (Vihervaara *et al*., 2017). Instead, the predominant function of HSF1 in activating eRNA transcription appeared to involve the recruitment of Pol II. First, eRNA synthesis was minute along the whole length of HSF1-dependent enhancers in the absence of HSF1 (Figure 5B). Second, Pol II did not accumulate at enhancer pause regions in HSF1-knockdown cells (Figure 5B-C,E; Figure S4D), which would be expected if HSF1 primarily released the pause at enhancers. The HSF1-assisted Pol II recruitment could be indirect, mediated for example by chromatin modifications that follow HSF1-binding (Zobeck *et al*., 2010; Petesch and Lis, 2012; Kusch *et al*., 2014; reviewed in Vihervaara and Sistonen, 2014). Alternatively, promoter-proximal pause-release has been shown to drive new initiation (Gressel *et al*., 2017), which might bring the transcription machinery also to functionally connected enhancers. In preconditioned cells, the heat-induced recruitment of Pol II to HSF1-dependent enhancers was diminished (Figure 5B; Figure S4D), which resembled the reduced initiation detected at heat-induced promoters (Figure 3F; Figure 4B-C). This globally reduced initiation in the presence of a potent *trans*-activator points to a systemic, HSF1-independent decrease in Pol II availability in preconditioned cells.

### Pol II Decelerates at CPSs of Actively Transcribed Genes

To monitor distribution of the transcription machinery upon heat shock in unconditioned *versus* preconditioned cells, we counted Pol II molecules that were engaged in transcription of enhancers, divergent transcripts, promoter-proximal regions, gene bodies, CPSs, and regions downstream of CPSs (Figure S5A-B). As expected, Pol II engagement was equal along the gene body regions of heat-induced genes in unconditioned and preconditioned cells, and its engagement was lower at heat-activated regulatory elements after preconditioning (Figure S5B). This decreased Pol II engagement at genes and enhancers occurred concurrently with increased Pol II occupancy downstream of the CPS, both at heat-induced and heat-repressed genes (Figure S5B). Since heat-repressed genes also showed increased Pol II density downstream of the CPS, we searched for genes with simultaneously reduced 5’ and increased 3’ density in heat-shocked preconditioned cells. The identified 415 genes (Figure 6A) comprised almost exclusively of genes that either gained (heat-induced genes) or retained (unchanged or heat-repressed genes) high transcription upon heat shock (Figure 6B-C, Figure S5C), which coupled the prolonged 3’ residency in preconditioned cells to high gene activity. The increased Pol II density in preconditioned cells was confined to 4 kb downstream of the CPS (Figure 4C; Figure 6A), demonstrating that Pol II residence time increased at the region where nascent transcript is cleaved, pre-mRNA polyadenylated, and transcription terminated. This decelerated Pol II progression through the CPS remarkably differs from previously reported stress-induced run-through transcription that can extend tens of kilobases downstream of the CPS (median 8.9 kb), and does not locally confine Pol II to the CPS (Vilborg *et al*., 2017; exemplified in Figure S3E).

**Figure 6.**
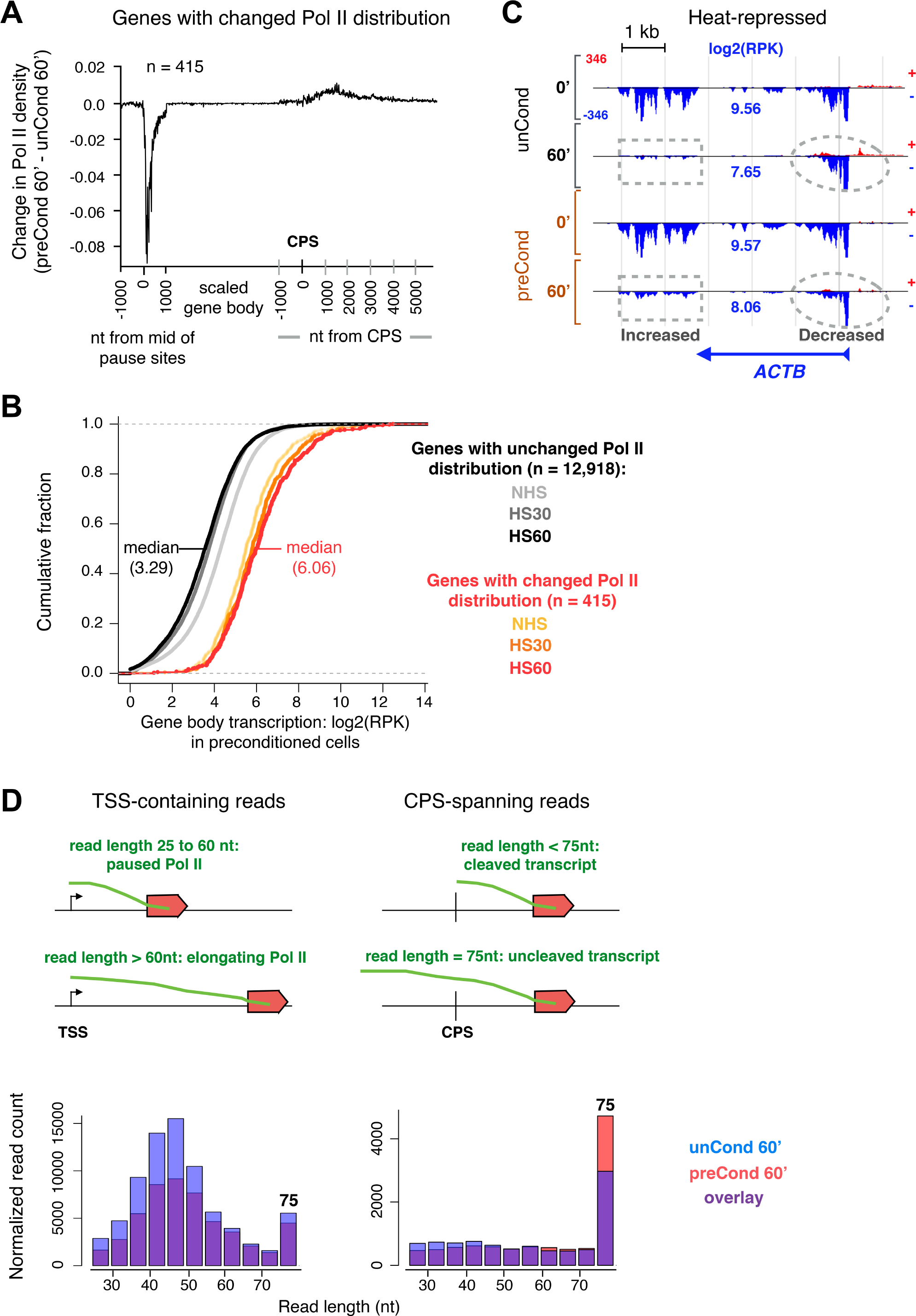
Pol II deceleration at CPSs co-occurs with reduced transcript cleavage and decreased genome-wide initiation. **A)** Difference in heat-induced Pol II density between preconditioned and unconditioned K562 cells, as measured along genes with reduced Pol II engagement at the 5’-end and increased engagement downstream of the CPS. **B)** Gene body transcription (log2RPK) was plotted cumulatively for genes with unchanged (shades of gray) or changed (shades of orange) distribution of Pol II due to preconditioning. Genes with changed Pol II distribution constitute highly transcribed genes (median log2 RPK: 6.06), whereas genes with no detectable change (shades of gray) show modest transcription level (median log2RPK: 3.29). **C)** Heat-repressed *beta-actin (ACTB)* displays reduced Pol II engagement at the 5’ region (dashed circles), and increased engagement downstream of the CPS (dashed squares) in preconditioned *versus* unconditioned cells upon heat shock. Despite prominent heat-repression, the gene body transcription of *ACTB* (log2RPK in blue) remains high (8.06) upon heat shock. **D)** Upper left panel: schematic presentation of how the lengths of TSS-overlapping PRO-seq reads report the position from the TSS to which Pol II has transcribed. Note that the maximum read length in this study is 75 nt. Lower left panel: histogram of reads at promoter-proximal regions showing that the decrease in Pol II density in preconditioned *versus* unconditioned cells upon heat shock consists of transcripts associated with paused (25 to 60 nt reads) and early elongating (>60 nt reads) Pol II complexes. Upper right panel: schematic presentation of how reads downstream of the CPS report events of transcript cleavage. Lower right panel: Histogram of CPS-containing reads showing that the increase in Pol II density in preconditioned *versus* unconditioned cells upon heat shock consists ofuncleaved transcripts.

### Decelerated Termination Co-Occurs with Reduced Transcript Cleavage

The prolonged residence time of Pol II at the CPSs, together with decreased initiation at heat-activated genes and enhancers, suggested reduced recycling of the transcription machinery from the ends of active genes to heat-induced transcription. To examine the status of Pol II and processing of transcripts at 5’- and 3’-ends of genes, we analyzed the length distribution of PRO-seq reads that overlap either with the transcription start site (TSS) or CPS. In essence, Pol II that undergoes initiation or pausing has transcribed through fewer nucleotides (<60 nt) than the sequenced read length in our PRO-seq data (75 nt). Thereby, the TSS-containing reads report, besides the initiating base (5’-end of the read), also the active site of transcription (3’-end of the read) of each individual transcript (Figure 6D; Rasmussen *et al*. 1993; Nechaev *et al*., 2010; Tome *et al*., 2018). The lower density of promoter-proximal Pol II that was detected after heat shock in preconditioned cells (Figure 6A; Figure S5B), comprised of transcripts with the whole spectrum of PRO-seq read lengths (20 nt to 75 nt), which indicates less initiating, pausing and early elongating Pol II complexes (Figure 6D). This reduction in all promoter-proximal Pol II states can be explained by less Pol II molecules engaging into transcription at gene promoters.

At the region intimately downstream of the CPS, the read length provides a measure of transcript cleavage: Reads that are shorter than 75 nt represent transcripts that have been cleaved to release the pre-mRNA (Figure 6D). After the cleavage, the uncapped nascent transcript is targeted by 5’-3’ exoribonuclease that degrades the RNA until reaching Pol II and transcription terminates (reviewed by Proudfoot, 2009). Measuring the read lengths downstream of the CPS revealed that the genome-wide increase in Pol II density at the 3’-ends of genes (Figure 6A; Figure S5B) comprised almost exclusively of reads with the maximum length of 75 nt (Figure 6D). This selective increase in transcription complexes with no signs of cleavage indicated that the decelerated progression of Pol II at the CPS co-occurred with reduced RNA processing. Taken together, the extended residence time of Pol II at the CPS, the reduced cleavage of the associated RNA, and the diminished initiation at heat-induced genes and enhancers provided evidence for genome-wide decrease in recycling Pol II from the ends of active genes to heat-induced regulatory elements.

## Discussion

### Control of Pol II Pause-Release Enables Rapid and Reversible Transcriptional Reprogramming

The groundbreaking model by Conrad Waddington describes developing cells as marbles that roll down an energy landscape of hills and valleys. While rolling down, cells take different paths and commit to distinct cell types, remodeling their chromatin environment and changing their transcription program (Waddington, 1957; reviewed in Takahashi and Yamanaka, 2015). Reversing from a differentiated to pluripotent cell, instead, requires specific transcription factors that push the cell up the energy landscape, which rarely occurs in nature (Gurdon *et al*., 1958; Takahashi and Yamanaka 2006). In this study, we showed that after genome-wide reprogramming of transcription by heat shock, cells robustly return to their cell type-specific transcription program within hours of recovery (Figure 1). In Waddington’s landscape, the heat-induced reprogramming of thousands of genes would be analogous with the cell transiently occupying a near-by valley, but during recovery, returning to its cell type-specific basal transcription program without acquiring new functions or committing to differentiation. This rapidly reprogrammed and precisely recovered transcription of genes and enhancers highlights the plasticity of transcription program, and implies that the transcriptional heat shock response is truly transient.

The ability of a cell to rapidly and robustly restore its cell type-specific transcription after heat shock is mechanistically explained by the genome-wide control of promoter-proximal Pol II pause-release. An important consequence of repressing thousands of genes by preventing the release of Pol II from their promoter-proximal regions is its rapid reversibility; a simple re-activation of the pause-release can restore productive gene transcription genome-wide without extensive chromatin remodeling. In this regard, Pol II pausing can be considered as one form of a memory that marks active genes during their transient repression. Indeed, while reprogramming of transcription during differentiation involves gene silencing and activation by remodeling the chromatin (reviewed in Perino and Veenstra, 2016), the reported changes in chromatin state upon heat shock comprise parylation of activated genes (Zobeck *et al*., 2010; Petesch and Lis, 2012), sumoylation of promoter-proximal regions (Niskanen *et al*., 2015), and increased histone acetylation and nucleosome accessibility at the sites of increased Pol II engagement, including the pause-region of repressed genes (Mueller *et al*, 2017; Vihervaara *et al*, 2017). These modifications are likely transiently compartmentalizing the transcription machinery and distinct gene activities in heat-shocked cells (reviewed in Vihervaara *et al*., 2018). Moreover, chromatin conformation, as measured by Hi-C, remains stable upon heat shock (Ray *et al*., 2019), which implies that the rapid recovery from stress does not require re-wiring of the chromatin connectivity. We, therefore, conclude that as chromatin architecture is primed for an instant transcriptional response to heat shock (Vihervaara *et al*., 2017; Ray *et al*., 2019), the Pol II pausing at heat-repressed genes primes rapid and robust transcriptional recovery.

### Stress-Induced Control of Pol II Is Carried over Mitotic Divisions

Tracking nascent transcription control that underlies transcriptional memory further emphasizes the importance of Pol II pause-regulation in genome-wide stress responses. A single heat shock, which is unlikely to cause permanent or long-lasting damage to the cell, did not change the basal transcription, but it increased Pol II pausing (Figure 1). Moreover, a single heat shock accelerated heat-induced transcription in the daughter cells by allowing more efficient progression of Pol II through the pause-region (Figure 2). This faster responsiveness without a change in the basal transcription mechanistically explains how cells can retain a memory of previously encountered stress to prepare their daughter cells for protection against proteotoxicity (Figure 7).

**Figure 7.**
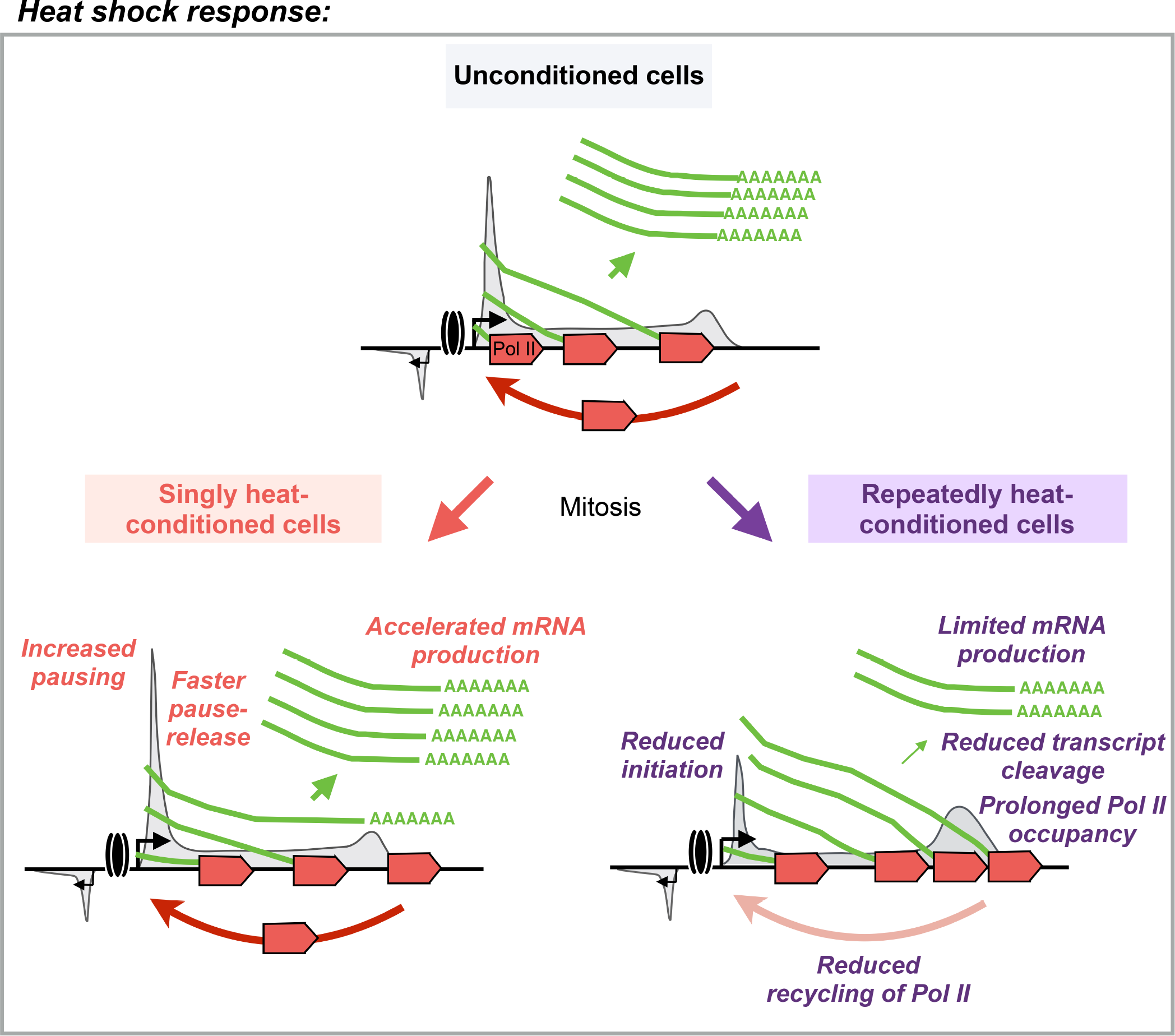
Heat-induced transcriptional memory is carried over mitotic divisions to accelerate promoter-proximal pause-release and decelerate termination. In unconditioned cells (upper panel), paused Pol II is rapidly released from the promoters of heat-induced genes into elongation, and it efficiently proceeds through the gene. A single heat shock exposure (lower left panel) primes an additional set of genes for instant heat-induction in the daughter cells by increasing Pol II pausing and by triggering a more rapid release of Pol II into productive elongation. Multiple heat shocks (lower right panel) cause reduced transcript cleavage at the 3’-end of active genes, which decelerates termination, and decreases recycling of the transcription machinery to heat-activated genes and enhancers.

Hallmarks of cancer cells include high stress-tolerance and active proliferation even in the presence of sustained stress (Hanahan and Weinberg, 2011). In accordance, human K562 erythroleukemia cells proliferated through multiple heat shocks that challenge the proteome integrity (Figure S3). The persistence of protein-damaging stress in the daughters of repeatedly stressed K562 cells was reflected in their transcriptional profile; two mitotic divisions after nine heat exposures, transcription of certain pro-survival genes was elevated, whereas expression of genes that are involved in maintaining protein production was decreased (Figure 3). The declined synthesis of protein production machinery was accompanied with reduced processing of transcripts at the 3’-ends of active genes (Figure 6). In cells with decreased protein synthesis, the decelerated transcription termination likely serves to reduce the mRNA load, as less pre-mRNA is released from the chromatin, and fewer Pol II molecules become available for new rounds of heat-induced transcription (Figure 7). Moreover, increased association of uncleaved transcripts at the 3’-ends of genes could provide a reservoir of pre-mRNAs that are rapidly processed to mature mRNAs once proteotoxicity is relieved and the cell restores its protein synthesis. This regulatory step at the 3’-end of the gene would be particularly important to instantly restore production of mRNAs from long genes, where transcription, proceeding on average 2.5 kb/minute (Jonkers *et al*., 2014), can take hours to complete after the Pol II pause-release. Intriguingly, the mechanisms that primed gene activation and refined transcription over mitotic divisions were mediated *via* regulation of Pol II, but they were not detected to involve altered activity of HSF1 (Figure 4; Figure 7). Taken together, in cells exposed to either single or multiple heat shocks, the re-wiring of transcriptional responsiveness mechanistically uncovered how regulation of Pol II can be passed over mitotic cell divisions to change the daughter cells’ response to external stress stimuli. These results revealed that experienced conditions can prime transcriptional responsiveness in the daughter cells, and provided evidence for a mechanistic control of nascent transcription machinery that maintains a memory over mitotic divisions.

## Materials and Methods

### Cell Culture, Heat Treatments and Cell Cycle Profiling

Cells were maintained at 37°C in a humidified 5% CO_2_ atmosphere. MEFs (McMillan et al., 1998) were cultured in Dulbecco’s modified medium (Gibco), and K562 cells in RPMI (Sigma), supplemented with 10% FCS, 2 mM L-glutamate, and streptomycin/penicillin. Cells were heat treated at 42°C either by submerging the cell culture into a water bath (K562 cells) or placing in an incubator after instantly provoking the heat shock with pre-warmed pre-conditioned media (MEFs). Recovery from the heat shock(s) was conducted at 37°C in a humidified 5% CO_2_ atmosphere. DNA content of the cells was determined by propidium iodide (PI) staining (40 μg/mL; Sigma), and progression of the cell cycle monitored by fluorescence-mediated counting (FACSCalibur, BD Biosciences). The FACS histograms were generated using Cell Quest Pro-6.0 (BD Biosciences) and Flowing Software 2.5 (Turku Centre for Biotechnology). The error bars in statistical analyses indicate standard deviations.

### Depletion of HSF1 with Short Hairpin RNA

HSF1 was depleted from K562 cells as previously described (Östling *et al*., 2007; Vihervaara *et al*., 2013) using shRNA constructs ligated into pSUPER vectors (Oligoengine). The vector-encoded oligonucleotides recognized HSF1 mRNA (GCT CAT TCA GTT CCT GAT C), or contained a scrambled sequence (GCG CGC TTT GTA GGA TTC G) that is not predicted to bind any sequence encoded by the human genome. The shRNA constructs were transfected into cells by electroporation (970 μF, 220 mV) 24 hours prior to the first heat treatment.

### Quantitative Reverse Transcription PCR

For analyzing polyadenylated mRNA, RNA over 200 nt was first isolated using RNeasy kit (Qiagen). Subsequently, 1 μg of RNA was treated with DNase I (Promega) and reverse transcribed with Moloney murine leukemia virus reverse transcriptase RNase H(−) (Promega) using oligoT primer. Quantitative PCR (qPCR) reactions were run using ABI Prism 7900 (Applied Biosystems) with primers (Oligomer) and probes (Oligomer or Roche Applied Science) reported in Vihervaara *et al*. (2013) and Elsing *et al*. (2014). HSP mRNA levels were normalized to mRNA of *GAPDH*, and fold inductions calculated against non-treated (unCond 0’) cells. All reactions were made in triplicate for samples derived from at least three biological replicates. Standard deviations were calculated and are shown in the graphs.

### Western Blotting

Cells were lysed in buffer C (25% glycerol, 20 mM Hepes pH 7.4, 1.5 mM MgCl_2_, 0.42 M NaCl, 0.2 mM EDTA, 0.5 mM PMSF, 0.5 mM DTT), and protein concentration in the soluble fraction was measured using Bradford analysis. Twenty micrograms of total soluble protein was boiled in Laemmli sample buffer, subjected to SDS-PAGE and transferred to nitrocellulose membrane (Protran nitrocellulose; Schleicher & Schuell). Proteins were analyzed with primary antibodies against HSF1 (Spa-901; Enzo) and β-tubulin (ab6046; Abcam). The secondary antibodies were HRP conjugated (GE Healthcare), and the blots were developed using an enhanced chemiluminescence method (ECL kit; GE Healthcare).

### PRO-seq

PRO-seq was performed as previously described (Kwak *et al*., 2013; Mahat *et al*., 2016; Vihervaara *et al*., 2017) with minor modifications. Nuclei of K562 cells were isolated in buffer A (10 mM Tris-Cl pH 8.0, 300 mM sucrose, 3 mM CaCl_2_, 2 mM MgAc_2_, 0.1% TritonX-100, 0.5 mM DTT) using Wheaton homogeniser (#357546, loose pestle). MEFs were incubated in permeabilization buffer (10 mM Tris-Cl, pH 7.5, 10 mM KCl, 250 mM sucrose, 5 mM MgCl_2_, 1 mM EGTA, 0.05% Tween-20, 0.5 mM DTT, 1x protease inhibitors [Roche], 0.4 u/μl RNase inhibitor). The nuclei or permeabilized cells were flash-frozen and stored at −80°C (10 mM Tris-HCl pH 8.0, 25% glycerol, 5 mM MgAc_2_, 0.1 mM EDTA, 5 mM DTT). Equal amount of untreated *Drosophila* S2 cells was spiked into each sample, and the run-on reaction performed at 37°C for 3 minute in the presence of biotinylated nucleotides (5 mM Tris-HCl pH 8.0, 150 mM KCl, 0.5% Sarkosyl, 2.5 mM MgCl_2_, 0.5 mM DTT, 0.05#mM biotin-A/C/G/UTP [Perkin Elmer], 0.4 u/μl RNase inhibitor). The total RNA was isolated with Trizol LS (Invitrogen). After EtOH-precipitation, the RNA was air-dried, base hydrolyzed with 0.1 N NaOH for 5 minutes on ice, and the NaOH was neutralized with Tris-HCl (pH 6.8). Unincorporated nucleotides were removed using P-30 columns (Bio-Rad), and the biotinylated nascent transcripts were isolated in a total of three rounds of streptavidin-coated magnetic bead (M-280, Invitrogen) purifications. Each of the bead binding was followed by Trizol extraction and EtOH-precipitation of the transcripts. The 5′-cap was removed with RNA 5’ Pyrophosphohydrolase (Rpph, NEB), and the 5′-hydroxyl group was repaired with T4 polynucleotide kinase (BioLabs). The libraries were generated using TruSeq small-RNA adaptors and sequenced using NextSeq500 (Illumina).

### PRO-qPCR

To quantify nascent RNA synthesis from selected heat-responsive genes, we modified PRO-seq to perform qPCR after the 3’-adaptor ligation. In brief, run-on reactions were conducted in the presence of both unlabeled (200 μM A/C/G/UTP) and biotinylated (50 μM biotin-A/C/G/UTP) nucleotides during a 5-minute run-on reaction at 37°C. Total RNA isolation, base hydrolysis, and 3’ adaptor ligation were conducted as described for PRO-seq. After the second bead binding, reverse transcription was performed using a primer against the 3’ adaptor, and qPCR reactions run with ABI Prism 7900 (Applied Biosystems). Primers (Oligomer) and probes (Oligomer and Roche Applied Sciences) were: *HSPH1* forward: AGCAGGCGGATTGTTGTTAG; *HSPH1* reverse: AAAGAGGTGGGCTAATCTTTCA; *HSPH1* probe: #38 (universal probe library, Roche); *HSPA1A* forward: GCCGAGAAGGACGAGTTTGA; *HSPA1A* reverse: CCTGGTACAGTCCGCTGATGA; *HSPA1A* probe: FAM-TTACACACCTGCTCCAGCTCCTTCCTCTT-BHQ1; *MED26* forward: ATTCCAGATGACCCGCTAAG; MED26 reverse: CGGATCACTACCACACCAGA; *MED26* probe: #21 (universal probe library, Roche). The nascent transcription of *HSPA1A* and *HSPH1* was compared against nascent transcription of *Mediator subunit 26* (*MED26*), a gene and a region in the gene that was actively transcribed and unchanged upon heat shock (Vihervaara *et al*., 2017).

### Computational Analyses of PRO-seq Data

The PRO-seq reads were adapter-clipped using cutadapt (Martin, 2011) and trimmed and filtered with fastx toolkit (http://hannonlab.cshl.edu/fastx_toolkit/). Reads from K562 cells were aligned to the human genome (GRCh37/hg19), and reads from MEFs to the mouse genome (mm10), using Bowtie 2 (Langmead and Salzberg, 2012). Unless otherwise stated, Bowtie 2 was run with standard parameters and only uniquely mapping reads were retained. The complete raw data in K562 cells (GSE127844), and MEFs (GSExxxxx) is available through Gene Expression Omnibus database (https://www.ncbi.nlm.nih.gov/geo/).

### Normalization of PRO-seq Data

Since heat shock provokes genome-wide changes in transcription and our samples were subjected to several days of treatments and recovery, an external normalization control was required to ensure correct PRO-seq sample comparison. To generate an invariant normalization vector in each sample, we spiked-in the same amount of non-treated *Drosophila* S2 cells (isolated nuclei or permeabilized cells) to each run-on reaction. Since run-on reaction adds biotinylated nucleotides to transcripts regardless of the species, this spike-in material follows through the biochemical and data-analytical steps of PRO-seq. Consequently, the spike-in material provides a read-out of uniquely mapped reads to the *Drosophila* genome, against which the sample data can be normalized. When mapping the data, we combined the sample genome with the *Drosophila* genome (hg19-dm3 or mm10-dm3), and retained the reads that uniquely mapped to the sample genome (hg19 or mm10) or the spike-in genome (dm3).

Normalization with spike-in reads demonstrated highly similar transcription profiles in unconditioned and preconditioned cells (Figure S1B; Figure S3C). This allowed us to utilize also samples generated in our previous study (Mahat *et al*., 2016; GSE71708) that highly correlated (rho=0.96) with the data generated in this study (Figure S1H). Since our earlier data did not have spike-in reads we utilized the ends of long genes when normalizing samples between distinct datasets. As described in Mahat *et al*. (2016), the ends of long genes provide an invariant region of transcription where the advancing or receding wave has not reached during a limited heat-shock time course.

### Quantifying Gene Transcription

To identify actively transcribed genes and their primarily used isoforms, transcription initiation sites were mapped genome-wide using discriminative regulatory elements identification from global run-on data (dREG; *https://dreg.dnasequence.org*). In essence, dREG gateway is trained to call transcription initiation sites of genes and enhancers (Wang *et al*., 2019) using their characteristic pattern of transcription (Core *et al*., 2014; Tome *et al*., 2018). To identify gene isoforms with active transcription initiation, TSSs of RefSeq-annotated transcripts were intersected (Quinlan and Hall, 2010) with dREG-called active regulatory elements. Subsequently, transcripts that harbored dREG-called initiation at the TSS were retained. The level of transcription *per* each transcript was measured from the gene body (+500 nt from TSS to −500 nt from CPS), as described previously (Mahat *et al*., 2016; Vihervaara *et al*., 2017). In the downstream analyses, we retained a single transcript *per* gene by selecting the isoform that showed the largest fold change to heat shock, or if called unresponsive to heat stress, had the highest level of transcription in non-stress condition. The analyses of enriched gene annotation categories were performed with Database for Annotation, Visualization and Integrated Discovery (DAVID; Dennis *et al*., 2003).

### Identification of Transcribed Enhancers

Nascent transcription architecture can be used for genome-wide *de novo* identification of active enhancers (Azofeifa *et al*., 2017; Vihervaara *et al*., 2017; Chu *et al*., 2018; Henriques *et al*., 2018; Wang *et al*., 2019). To call transcribed enhancers across the genome, dREG gateway (https://dreg.dnasequence.org/) was utilized to identify initiation regions from the PRO-seq datasets. Subsequently, initiation regions that did not overlap with the TSS of any annotated transcript (RefSeq) were retained. This class of distal regulatory elements initiates transcription and includes transcriptionally active enhancers. Enhancer transcription modestly correlates with functional enhancer activity (Henriques *et al*., 2018; Michaylichenko *et al*., 2018; Tippens *et al*., 2018). However, we wish to emphasize that eRNA production does not provide a direct read-out for enhancer’s ability to activate transcription of distally located regulatory element(s). Because heat shock can change the pattern of Pol II progression at enhancers (Vihervaara *et al*., 2017), we first identified putative enhancers individually in each sample, and then unified the enhancer coordinates between samples using bedtools merge with d −100 (Quinlan and Hall, 2010).

### Analyses of Differential Gene and Enhancer Transcription

To call significant changes in gene and enhancer transcription, we utilized DESeq2 (Love *et al*., 2014), which uses the variance in biological replicates to assess significant changes between conditions. Differential gene expression was quantified using gene body transcription, whereas enhancer transcription was quantified along the whole enhancer length, individually for minus and plus strands (Vihervaara *et al*., 2017). For significantly changed transcription, we required p-value <0.05 (K562) or <0.001 (MEFs), and fold enrichment >1.25. The less stringent criterion for K562 cells used in this study, as compared to MEFs and our earlier data on K562 cells (Vihervaara *et al*., 2017), is due to lower sequencing depth. The identified genes and enhancers are highly similar in distinct studies of the same cell type.

### Analyses of HSF1-Dependent Transcription at Genes and Enhancers

HSF1-knockdown data was inferred from a single replicate, chosen by the most prominent down-regulation of HSF1 throughout the length of the experiment (Figure 5A). To identify HSF1-dependent genes, we used two approaches. First, we measured the heat-induced gene body transcription for each gene in the presence and absence of HSF1. This comparison of transcription level identified 186 genes whose heat induction in K562 cells depleted of HSF1 remained under 50% of the respective induction in cells expressing intact HSF1. Second, we used the fact that unconditioned and preconditioned cells correlated to the same extent as biological replicates (rho=0.98) and contained similar levels of gene body transcription (Figure S5B), conducting DESeq2 using the same time point from unconditioned and preconditioned cells as a replicate pair. These analyses showed 227 genes and 496 enhancers to be HSF1 dependent in both unconditioned and preconditioned K562 cells (Figure S4B). Since a subset of genes changed basal or heat-inducible transcription due to preconditioning, we complemented the DESeq2-analysis by calling genes that showed HSF1-dependency only in unconditioned or preconditioned cells, or that were called insignificant due to changes in basal transcription. Since HSF1-dependency of the 227 DESeq2-called genes ranged from 64.1% to 99.9% (Figure S4B), we queried genes that in either unconditioned or preconditioned cells were HSF1-dependent at least to 64.1%, gained at least two-fold heat-induction, and had a minimum gene body transcription of 50 RPK in any condition. This analysis identified 18 additional HSF1-dependent genes, including *PPP1R15A* that had lost heat-inducibility, and *HSPA8* that had gained higher basal transcription, upon preconditioning. All of the 18 genes were individually verified to be HSF1-dependent by browsing.

### Visualizing Transcriptionally Engaged Pol II in Genome Browsers and as Composite Profiles

Pol II densities as bigWig and bedgraph files were visualized with Integrative Genomics Viewer (IGV; Thorvaldsdóttir *et al*., 2013) and an in-house browser (Hojoong Kwak, Cornell University, Ithaca, NY, USA). The read counts in defined genomic regions were obtained, and composite profiles generated using bigWig package (https://github.com/andrelmartins/bigWig/). The average intensities in composite profiles were queried in 20-nt, 10-nt or 1-nt bins. The shaded areas display 12.5-87.5% fractions of the data in each queried window. To generate an average profile of gene bodies with different lengths, 1/500th of the gene body length was set to the bin size, after filtering out short genes where the bin would have been less than 1 nt.

### Identification of Genes with Compromized Pol II Progression

To identify genes with decreased Pol II density in 5’- and increased density in 3’-region, genes were first divided into three distinct regions: 1) 5’-coding region comprising 1000 nt downstream of the mid coordinate between Pol II pause sites of divergent transcription, 2) gene body, measured from +1000 nt from the mid of the pause sites to −1000 nt from the CPS, and 3) downstream, +100 nt to +6000 nt, of the CPS. PRO-seq reads in each region were measured, after which the read count in the preconditioned 60-minute heat shock sample was deduced from the respective read count in the unconditioned 60-minute heat shock sample. Since gene body transcription varies from gene to gene, we compared the change in Pol II progression within each gene. To identify genes with reduced 5’- and increased 3’- Pol II density, we required the reduction at 5’-coding region to be three times larger than the absolute change in the gene body read count. Simultaneously, the increase in read counts downstream of the CPS was required to be three times higher than the absolute change in the gene body read count.

### Quantifying Engaged Pol II Molecules in Distinct Genomic Regions

The mapped reads were sorted to distinct genomic regions by intersecting the 3’-coordinate of the read with the genomic coordinates described in Supplementary Figure 5A. To avoid double mapping, gene body reads that overlapped with enhancers or pause regions were omitted. Subsequently, the number of reads in a given region was counted as fraction of total uniquely mapping reads in the PRO-seq data.

### Additional Datasets Used

Besides the PRO-seq data generated in this study (GSE127844 and GSExxxxx), the following datasets have been utilized: HSF1-binding sites in non-stressed and 30-minute heat-shocked K562 cells (GSE43579; Vihervaara *et al*., 2013), binding sites of TATA-box Binding Protein (TBP) in non-stressed K562 cells (GSM935495; Consortium EP, 2011); PRO-seq data in non-stressed and 30-minute heat-shocked K562 cells for verification purposes (GSE89230; Vihervaara *et al*., 2017); PRO-seq data in non-stressed and 12.5-minute heat-shocked MEF cells (GSE71708; Mahat *et al*., 2016).

### Code Availability

Computational analyses have been performed using Unix, R and Python languages. Custom made scripts can be made available upon request.

## Acknowledgements

We thank the members of the Sistonen and the Lis laboratories for valuable advice during the manuscript preparation. This work was financially supported by the Sigrid Jusélius Foundation (A.V., L.S.), Svenska Tekniska Vetenskapsakademin i Finland (A.V.), South-West Finland’s Cancer Foundation (A.V.), Joe, Tor and Pentti Borg Memory Foundation (A.V.), Åbo Akademi Research Foundation (A.V.), The Finnish Cultural Foundation (A.V.), Academy of Finland (L.S.), Cancer Foundation Finland (L.S.), Magnus Ehrnrooth Foundation (L.S.), Åbo Akademi University (L.S.), and NIH grant RO1-GM25232 (J.T.L.). The content is solely the responsibility of the authors and does not necessarily represent the official views of the National Institutes of Health.

## Supplementary Material

The supplementary material contains five figures and one dataset. The Supplementary Dataset 1 lists gene transcripts in human K562 cells that show transcription initiation at the TSS, identified from PRO-seq data using dREG gateway. The abbreviations in the Supplementary Dataset 1 are listed here. chr: chromosome. txStart: the first coordinate of the transcript (RefSeq). txEnd: the last coordinate of the transcript (RefSeq). Please note that txStart < txEnd, regardless whether the gene is on plus on minus strand. Strand: strand encoding the transcript. geneName: the name of the gene. txID: transcript specification. uC30_to_uC0: DESeq2-called regulation of transcription in unCond 30’ *versus* unCond 0’. pC0_to_uC0: DESeq2-called regulation of transcription in preCond 0’ *versus* unCond 0’. pC30_to_uC0: DESeq2-called regulation of transcription in preCond 30’ *versus* unCond 0’. UnExp: unexpressed genes (initiation of transcription is detected, but the level of engaged Pol II molecules on the gene body is very low). UnReg: unregulated. DownHC: down-regulated with high confidence (counted as down-regulated in this study). DownLC: down-regulated with low confidence. UpHC: up-regulated with high confidence (counted as up-regulated in this study). UpLC: up-regulated with low confidence. Please note that in the manuscript, only one transcript per gene is included in the downstream analyses.

**Supplementary Figure 1 (related to Figures 1 and 2).**
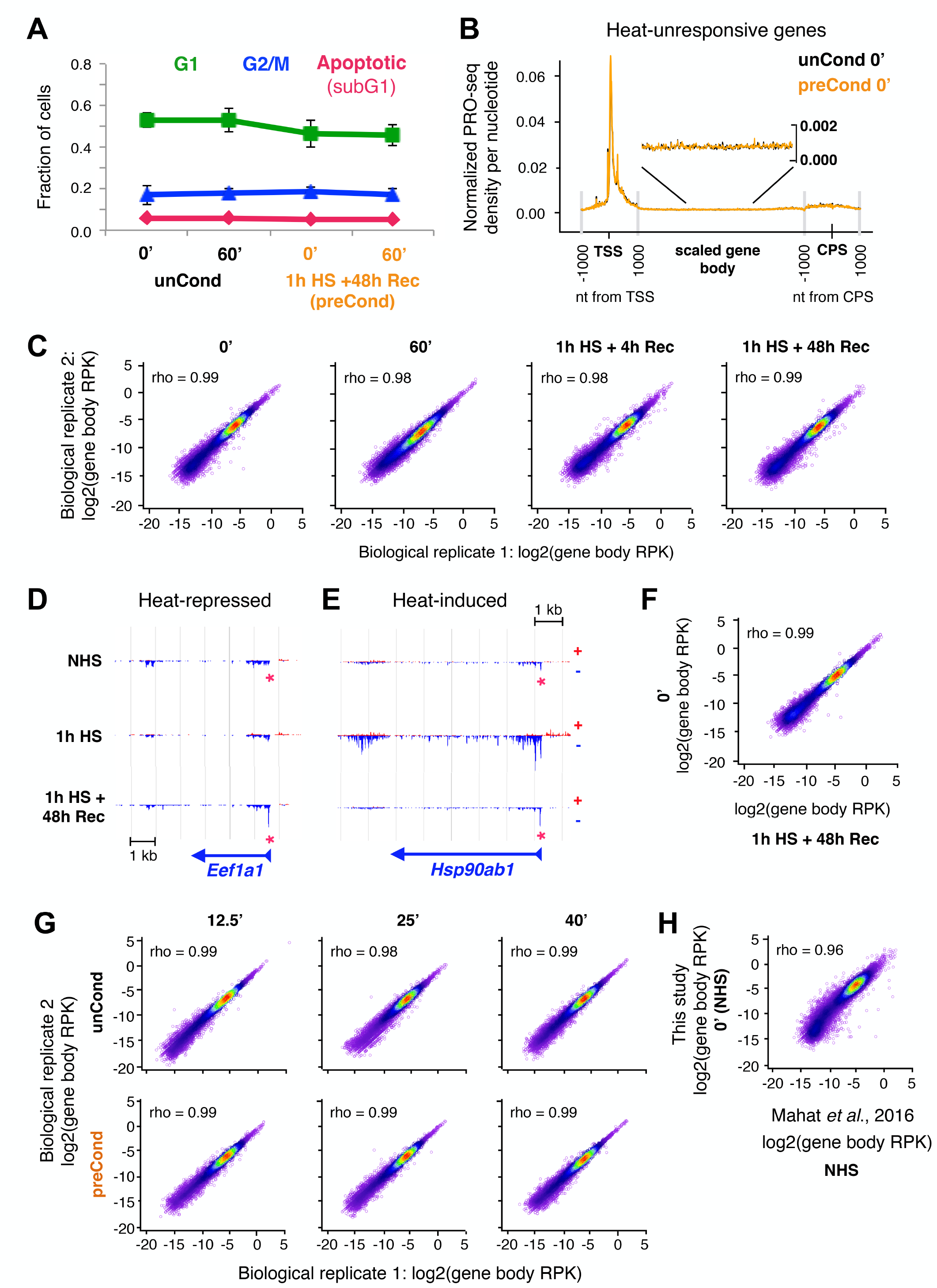
Whole-genome spike-in allows accurate data normalization and quantification of transcription responses. **A)** FACS-defined fraction of mouse embryonic fibroblasts (MEFs) with indicated DNA content during heat shock, recovery, and an additional heat shock exposure. Cells with a diploid genome are indicated with G1, and duplicated (tetraploid) genome with G2/M. Apoptotic refers to cells with fragmented genome (sub G1). **B)** Average profile of nascent transcription at heat-unresponsive genes in unconditioned and preconditioned MEFs after normalizing the datasets against whole genome run-on RNAs from spiked-in, permeabilized *Drosophila* S2 cells. **C)** Correlation plots of gene body transcription (log2RPK) between biological PRO-seq replicates. **D-E)** Transcriptional profile of a heat-repressed (D) and a heat-induced (E) gene, at which promoter-proximal Pol II pausing increases (D) or remains elevated (E) during recovery from a single heat shock. **F)** Correlation of gene body transcription (log2RPK) between non-heat-shocked cells (0’) and cells exposed to a 1-h heat shock alone or followed by a 48-h recovery (1h HS + 48h Rec). **G)** Correlation plots of gene body transcription (log2RPK) between biological PRO-seq replicates in unconditioned (upper panels) and preconditioned (lower panels) cells. **H)** Correlation plots of gene body transcription (log2RPK) between non-heat-shocked MEFs mapped in this study (y-axis) and in our previous study (x-axis; Mahat *et al*., 2016). In panels C, F-H, Rho indicates Spearman correlation.

**Supplementary Figure 2 (related to Figure 2).**
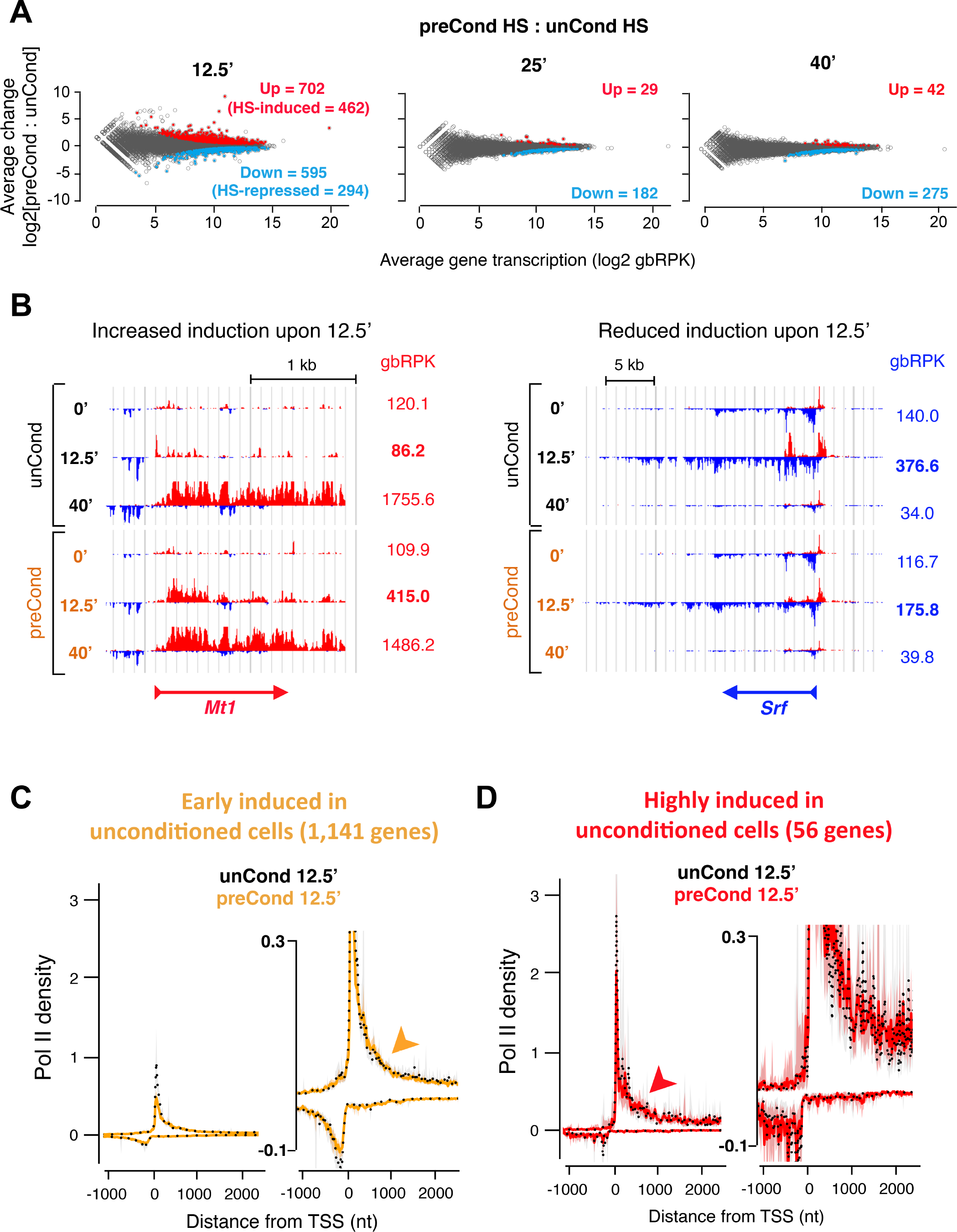
A subset of genes gains faster heat-responsiveness by preconditioning. **A)** DESeq2-analysis of differential gene transcription between unconditioned (unCond) and preconditioned (preCond) MEFs at the indicated time points of heat stress. Up denotes gene bodies with significant (p < 0.001) increase in heat-induced transcription in preconditioned cells as compared to unconditioned cells. Down denotes gene bodies where transcription in preconditioned cells is significantly (p < 0.001) lower than in unconditioned cells. In the 12.5-min time point, HS-induced indicates heat-induced genes that gain higher gene body transcription, and HS-repressed denotes heat-repressed genes that gain deeper transcription reduction, due to preconditioning. We refer to these genes as faster heat-induced and faster heat-repressed, respectively, throughout the manuscript. **B)** Browser shot examples of genes with increased (left panel) and reduced (right panel) heat induction in preconditioned cells. Gene body RPK (gbRPK) is indicated, and bolded in 12.5-min conditions that show statistically significant change between unconditioned and preconditioned cells. **C-D)** Average Pol II density at the promoter-proximal regions of genes that are called heat-induced upon the 12.5-min time point (C), or are highly heat-activated (D), in unconditioned cells. The Pol II density in C and D is compared between unconditioned (black dotted line) and preconditioned (yellow or red solid line) cells upon a 12.5-min heat shock. Arrowheads denote the signal downstream, of Pol II pausing, as in Figure 2D. The right panels in C and D depict the region around the pause in an expanded scale.

**Supplementary Figure 3 (related to Figure 3).**
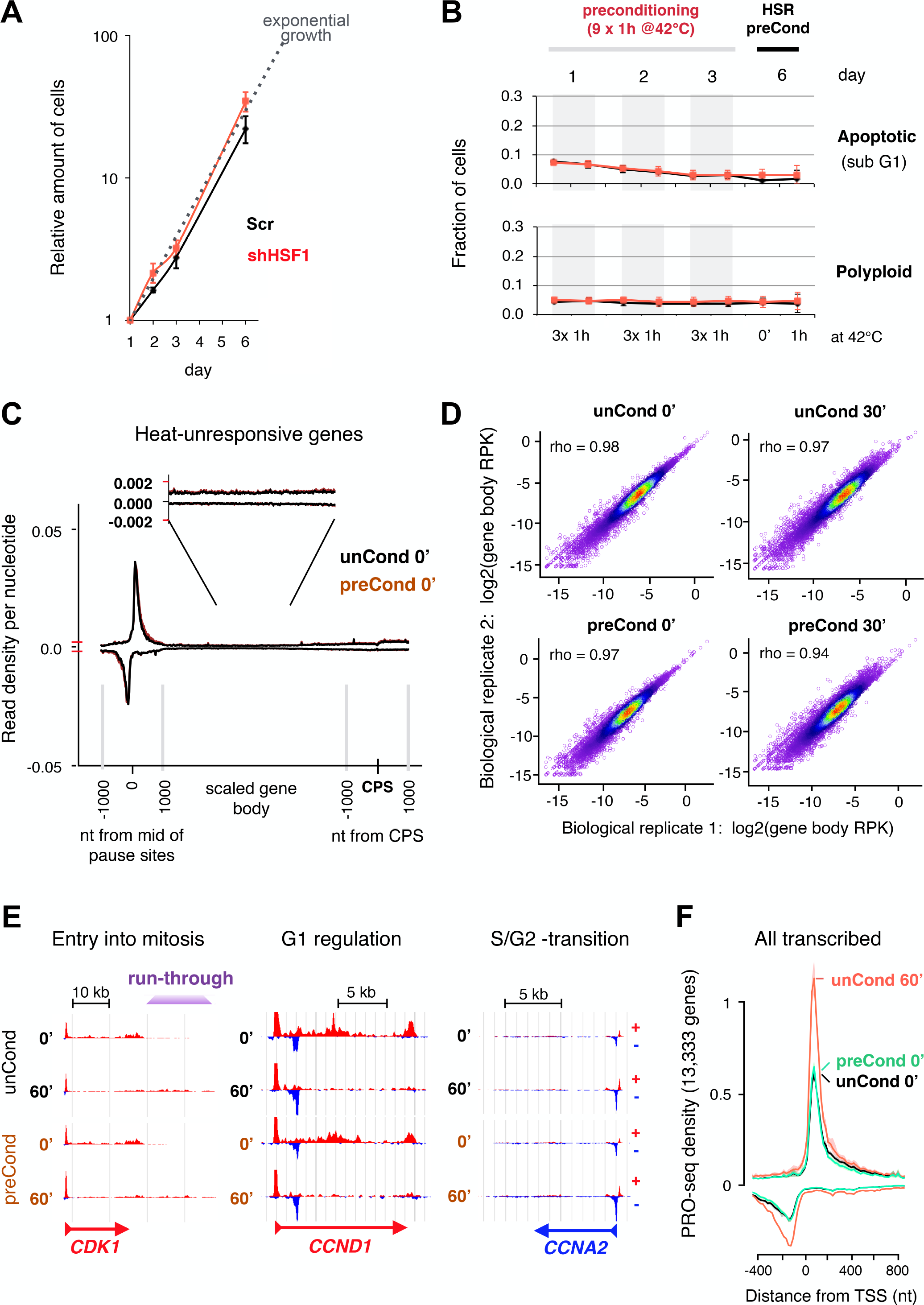
Human K562 cells proliferate through repeated stress conditions. **A)** Relative numbers of K562 cells in scrambled transfected (Scr) and HSF1-depleted (shHSF1) cells, as counted by dividing the number of cells at each day with the number of cells at day 1. The relative cell count is plotted in logarithmic scale, and theoretical exponential growth indicated with a dotted gray line, demonstrating that K562 cells maintain their proliferation rate throughout the preconditioning. **B)** FACS-defined fraction of cells with fragmented (indicative of apoptosis), or polyploid, genome. The slightly higher number of apoptotic cells in day 1 is likely a consequence of electroporation 24 h before the first heat shock treatment of preconditioning. **C)** Average Pol II density along the coding and non-coding strands of heat-unresponsive genes, shown in unconditioned and preconditioned cells after normalizing the samples against whole-genome spike-in from *Drosophila* S2 cells. Please note that the whole-genome (isolated nuclei) spike-in allows accurate normalization between the samples that were grown several days in distinct cell cultures. **D)** Correlation plots of gene body transcription (log2RPK) between biological PRO-seq replicates in unconditioned (upper panels) and preconditioned (lower panels) cells. Rho indicates Spearman correlation. **E)** Basal and heat-responsive transcription of cell cycle regulators *CDK1* (left panel), *CCND1* (middle panel), and *CCNA2* (right panel) in unconditioned and preconditioned K562 cells. Heat-induced run-through transcription (also known as Downstream of Genes, DoG) is indicated downstream of CPS in *CDK1*. **F)** Average Pol II density at the promoter-proximal regions of all transcribed genes. Please note that the average level of Pol II pausing is restored during the 48-h recovery from repeated heat shocks.

**Supplementary Figure 4 (related to Figures 4 and 5).**
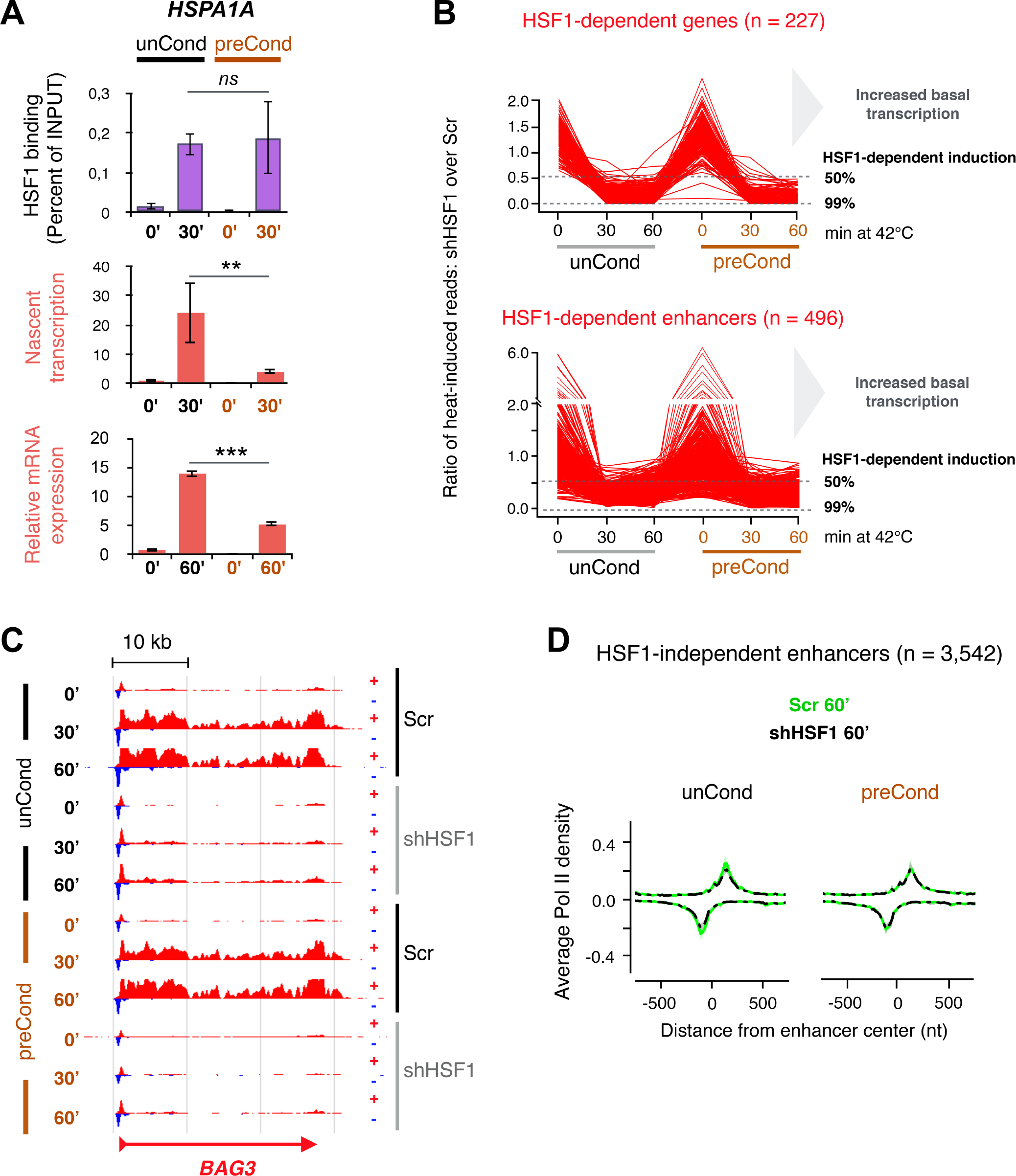
Repeatedly heat-shocked cells adjust transcription and mRNA expression over mitotic divisions. **A)** HSF1 binding at the promoter (uppermost panel), transcription of coding sequence (middle panel), and relative expression of polyA-containing mRNA (lowest panel) of *HSPAJA* in unconditioned and preconditioned K562 cells. ** indicates p-value < 0.05 and *** p-value < 0.005. **B)** HSF1­ dependent heat-inducibility of genes (upper panel) and enhancers (lower panel). HSF1-dependency for each individual gene or enhancer was measured by dividing transcription (gene body RPK) in shHSF1 by transcription in Ser, after basal transcription was removed from heat-treated samples. **C)** Transcription profile of HSF1-dependently heat­ induced *BAG3* gene in unconditioned and preconditioned cells in the presence (Ser) or absence (shHSF1) of HSF1. **D)** Average Pol II density at the HSF1-independently heat-induced enhancers in unconditioned and preconditioned cells in the presence or absence of HSF1.

**Supplementary Figure 5 (related to Figures 5 and 6).**
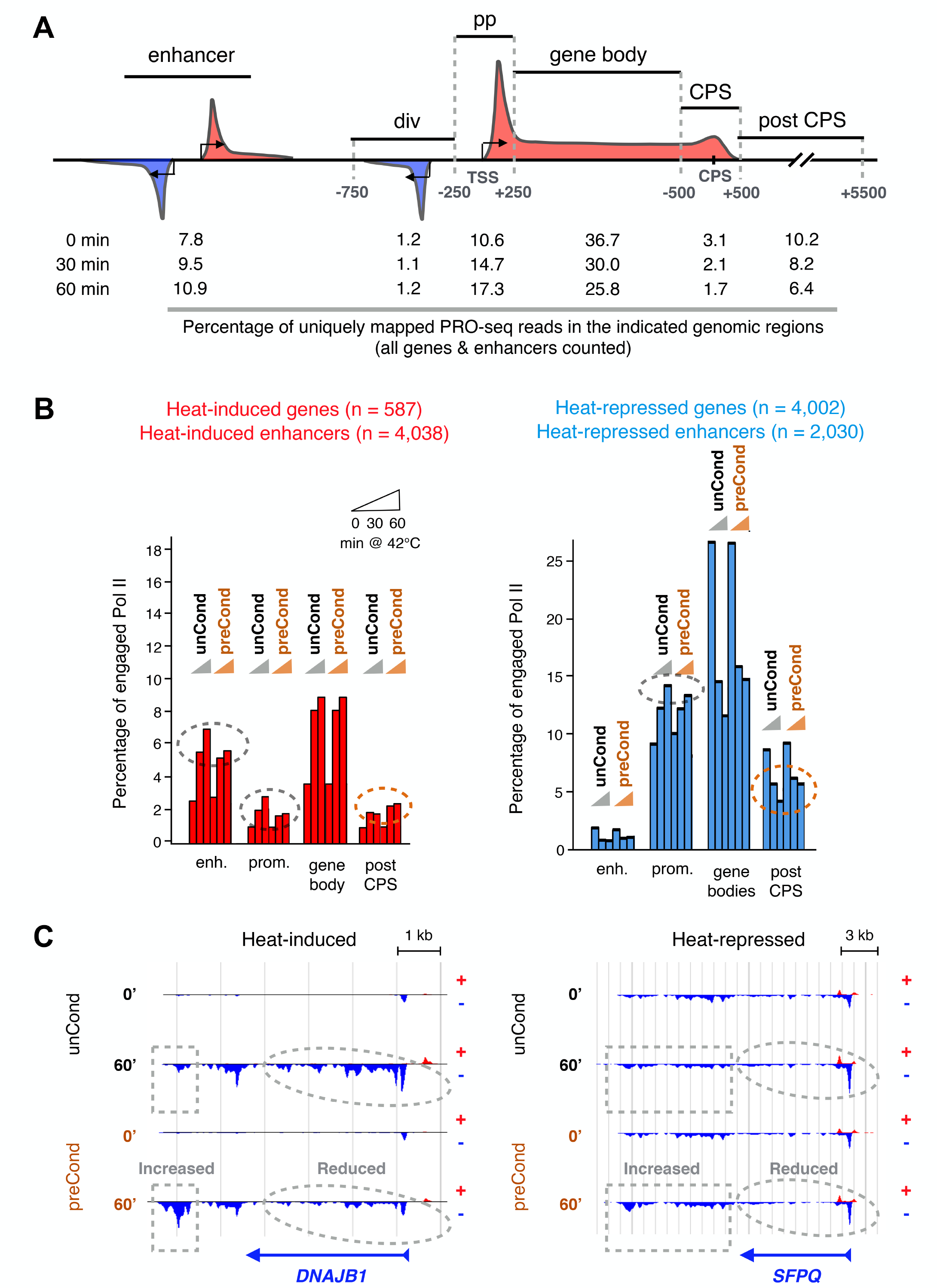
Genome-wide distribution of engaged Pol II molecules changes by preconditioning. **A)** Schematic representation of genomic regions within which the distribution of engaged Pol II molecules was measured. The numbers below each region indicate the percentage of all uniquely mapping PRO-seq reads that localize to the given region in unconditioned K562 cells. NHS: non-heat shock; 30 min and 60 min indicate the time of heat shock at 42°C. **B)** Percentage of all uniquely mapping PRO-seq reads at the indicated regions of heat-induced (left panel) and heat-repressed (right panel) genes and enhancers. Orange dashed circles indicate genomic regions with increased total engagement of Pol II in preconditioned cells. Gray dashed circles indicate regions with reduced Pol II engagement in preconditioned cells. The colored triangles above the bars denote the time of the heat shock. **C)** Transcription profile of heat-induced co-chaperone *DNAJB1* and heat-repressed splicing factor (*SFPQ*) genes. Dashed circles indicate regions with reduced, and squares with increased, Pol II engagement in heat-shocked preconditioned cells as compared to heat-shocked unconditioned cells.

